# Mitochondrial dysfunction underlies cardiac contractility and growth defects in a zebrafish model of *NAA15*-related heart disease

**DOI:** 10.1101/2025.08.04.668548

**Authors:** Warlen P. Piedade, Olivia Weeks, Hannah R. Moran, Alexander A. Akerberg, Michael M. Molnar, Jennifer Galdieri, Ashley Buick, Rongbin Zheng, Kaifu Chen, Hasmik Keshishian, Patrick Hart, Steven A. Carr, Calum A. MacRae, Caroline E. Burns, C. Geoffrey Burns

**Author notes:** Correspondence: C. Geoffrey Burns and Caroline E. Burns.

## Abstract

Delineating the genetic and environmental instigators of congenital heart disease (CHD), affecting up to 1% of newborns, will improve preventative, diagnostic, and therapeutic efforts to mitigate disease outcomes. Although mutations in *NAA15*, which encodes the auxiliary subunit of N-alpha-acetyltransferase A (NatA), are associated with CHD in humans, vertebrate animal models of *NAA15*-related heart disease have yet to be described. Therefore, we isolated zebrafish strains carrying null mutations in paralogs *naa15a* and *naa15b,* which we co-localized to the early larval myocardium. Double knockout (DKO) *naa15*-deficient larvae exhibited diminutive, lowly contractile, and bradycardic ventricles composed of fewer and smaller cardiomyocytes (CMs) incapable of proliferation. On a subcellular level, mutant CMs exhibited moderately disorganized myofibrils. CM-specific re-expression of *naa15a* partially rescued the contractility and growth deficits, revealing an indispensable myocardial function. Ubiquitous mis-expression of human *NAA15* achieved complete rescue, enabling functional testing of human *NAA15* variants. Animals with a reduced dosage (RD) of *naa15* survived to adulthood and shared phenotypes with DKO larvae, including smaller ventricular chambers and CMs, which exhibited mosaic myofibril disarray. Deep quantitative proteomic profiling of WT and *naa15^RD^* adult hearts revealed differential expression of multiple protein classes, including several subunits of mitochondrial respiratory complex I, a component of the electron transport chain. Accordingly, we documented reduced mitochondrial content and function in the myocardium of *naa15*-deficient larvae. Taken together, our data reveal myocardial functional and structural abnormalities associated with mitochondrial dysfunction in a vertebrate animal model of *NAA15*-related heart disease.

## Introduction

Among the most common and deadliest of congenital anomalies, congenital heart disease (CHD) affects ∼1% of births or ∼1.3M newborns annually worldwide^1,2^. Although mutations in ∼400 genes are predicted to cause CHD^3^, many genes remain unidentified. Moreover, for numerous currently recognized definitive and candidate CHD genes^4,5^, the pathogenetic mechanisms remain unknown or incompletely characterized, which impedes progress towards reducing CHD-associated morbidity and mortality.

The Pediatric Cardiac Genomics Consortium (PCGC) discovered that 3 CHD probands from 4511 trios were heterozygous for *de novo*, rare, and predicted loss-of-function (pLoF) variants in *N-alpha-acetyltransferase 15* (*NAA15*)^3,6–8^. Additionally, 15 probands were heterozygous for very rare inherited missense mutations. Because *NAA15* variants were more prevalent in the probands than in the general population (i.e., individuals cataloged in the Genome Aggregation Database; GnomAD), the PCGC concluded that *NAA15* haploinsufficiency likely contributes to CHD pathogenesis. Several CHD subtypes were observed across the 18 probands, including septal defects (atrial and ventricular), conotruncal malformations (Tetralogy of Fallot, double outlet right ventricle, transposition of the great arteries), and single ventricle disease (hypoplastic left or right heart syndromes; HLHS or HRHS). Despite the association between CHD and *NAA15* pLoF variants^6^, they are lowly penetrant and expressive^9^, revealing the influence of environmental, genetic, or epigenetic modifiers. In a separate study, damaging *NAA15* variants were identified in two patients with pediatric hypertrophic cardiomyopathy^10^. Beyond the heart, a convincing link has been drawn between damaging *NAA15* variants and neurodevelopmental disorders, including intellectual disability, autism spectrum disorder, and seizures, as well as dysmorphic facial features^6,9,11–15^. The full spectrum of abnormalities stemming from *NAA15* mutations is termed *NAA15*-related disorder or syndrome.

Together with the enzymatic subunit NAA10, NAA15 forms the core of the N-terminal acetyltransferase A (NatA) complex, which associates physically with the ribosome and acetylates the N-termini of target proteins co-translationally^16^. NatA-mediated acetylation occurs on Alanine, Cysteine, Glycine, Serine, Threonine, or Valine following cleavage of the initiator methionine and is a widespread protein modification, occurring on ∼40% of human proteins. Functionally, the N-terminal acetyl group increases local hydrophobicity, thereby influencing protein folding, stability, subcellular localization, and protein-protein interactions. NatA interacts with two other proteins, including huntingtin-interacting protein (HYPK), which promotes NatA activity, and NAA50, which is the catalytic subunit of NatE, of which Naa15 is also a subunit. On a molecular level, NAA15 tethers NatA to the ribosome and influences substrate specificity. Loss-of-function studies have demonstrated that *NAA15* is required for the viability of flies^17^, worms^18^, zebrafish^19^, and mice^15^. Mutant strains of budding or fission yeast were viable but exhibited growth defects^20–23^. Despite previous loss-of-function studies in zebrafish and mice, no information was provided on heart status.

A recent study analyzed human induced pluripotent stem cells (iPSCs) genetically engineered to carry *NAA15* pLoF mutations in heterozygous or homozygous states^6^. Heterozygous iPSCs were indistinguishable from wild-type (WT) cells. They differentiated into cardiomyocytes (iPSC-CMs) that exhibited normal contractility in 2D culture but reduced force production in 3D microtissues. By contrast, null iPSCs exhibited slow growth, low viability, and were incapable of directed CM differentiation, precluding phenotyping of null CMs. Proteomics analysis of iPSCs identified reduced N-terminal acetylation of a limited number of proteins in heterozygous (9) or null (32) iPSCs. It also identified differential expression of 562 proteins enriched for ribosomal subunits. Among the dysregulated proteins were four encoded by genes implicated in CHD pathogenesis, leading investigators to speculate that one or more are relevant downstream targets in *NAA15*-mediated CHD.

The association between NatA and pediatric heart disease extends beyond *NAA15*, as a LoF mutation in the enzymatic subunit gene *NAA10* was recently implicated in QT-prolongation and cardiomyopathy in a large family^24^. Recapitulating the variant in hiPSC-CMs revealed repolarization defects due to altered sodium and potassium currents, in accordance with QT prolongation. Moreover, variant-carrying engineered heart tissue exhibited sarcomere disorganization and reduced contractility, consistent with cardiomyopathy.

The embryonic function of zebrafish *naa15* was previously investigated using antisense morpholinos targeting paralogs *naa15a* and *naa15b*^19^. However, most double morphants died prior to heart development. Among the surviving animals, heart phenotypes were either absent or unreported, leaving open the question of how deleting or reducing *naa15* activity affects heart development. A *Drosophila* model of cardiac-specific knockdown of *naa15* was recently reported, but cardiac phenotyping was not possible because the animals failed to form a heart^25^.

In summary, although pLoF and missense *NAA15* variants have been linked to CHD and studied in *Drosophila* or human cell culture^3,6–8,10^, vertebrate animal models of *NAA15*-related heart disease are lacking. Filling this gap would complement invertebrate and human cell-based studies to explore pathogenic CHD mechanisms, test *NAA15* variants, and evaluate candidate therapies in pre-clinical studies^26,27^. To that end, we characterized larval hearts from *naa15*-deficient zebrafish, and adult hearts with reduced *naa15* dosage (*naa15^RD^*), revealing functional and structural myocardial abnormalities associated with mitochondrial dysfunction.

## Methods

### Zebrafish husbandry and strains

Zebrafish husbandry protocols^28^ were consistent with the Guide for the Care and Use of Laboratory Animals^29^ and were approved by the Institutional Animal Care and Use Committee (IACUC) of Boston Children’s Hospital. Live embryos, larvae, and adults were maintained at 28.5°C. The following zebrafish strains were utilized: wild-type (WT) TuAB, WT TD5, *TgBAC(-36nkx2.5:ZsYellow)^fb7,^*^30^, *Tg(myl7:nucGFP)^fb18^* [formerly *Tg(cmlc2:nucGFP)^fb18^*^,31^], *naa15a^chb13^* (this study), *naa15b^chb14^* (this study), and *Tg(myl7:nuc-mCherry-P2A-HA-naa15a)^chb15^*(this study).

### Brightfield, epifluorescence, and confocal microscopy

The following microscopes were used for imaging: a Leica fluorescence dissecting microscope with an Excelis AU-600-HDS HD Camera (Accu-Scope), a Nikon Eclipse 80i fluorescence compound microscope with a Retiga 2000R CCD camera (QImaging) controlled by NIS-Elements image acquisition and analysis software (Nikon Instruments), and an Olympus FLUOVIEW FV3000rs resonant scanning confocal microscope.

### NAA15 domain prediction and alignment

The UniProt^32^ protein sequences F1QE25 (*D. rerio* Naa15a), A0A0R4I9S5 (*D. rerio* Naa15b), and Q9BXJ9 (*H. sapiens* NAA15) were utilized for domain prediction by the InterPro database (https://www.ebi.ac.uk/interpro/)^33^ and alignment by Clustal Omega^34^. Generation, molecular characterization, and genotyping of fish carrying *naa15a^chb13^* and *naa15b^chb14^* mutant alleles Guide RNA targets in *naa15a* and *naa15b* were identified using CHOPCHOP^35^. Single guide RNAs targeting sequences 5’-GATGGTCTTGCCTCCATGTT<u>CGG</u>-3’ [<u>protospacer adjacent motif (PAM)</u>] in exon 2 and the downstream intron of *naa15a* (ENSDART00000017785; GRCz11) or 5’-AGGAGAGGAATCCGGAGAAC<u>TGG</u>-3’ in exon 7 of *naa15b* (ENSDART00000168067) were generated and microinjected with recombinant Cas9 protein (New England Biolabs) into one-cell stage zebrafish embryos, as described^36^. Founder fish with germline transmission of insertions or deletions (indels) to F1 offspring were identified using fluorescent PCR and DNA fragment analysis with primer pairs *15a-geno-F/R* and *15b-geno-F*/*R* (Table S1), as described^37^. Indel identities were determined through next-generation sequencing of mutant amplicons. The above primers were also used for genotyping animals. The WT *naa15a* and mutant *naa15b^ch13^* alleles produced amplicons of 199bp and 187bp, respectively, and the WT *naa15b* and mutant *naa15b^ch14^* alleles produced amplicons of 205bp and 210bp, respectively.

The major *naa15a* mRNA isoforms expressed in DKO animals were identified through PCR amplification of DKO cDNA with primer pair *15a-iso-F/R* (Table S1), which bound to exon 1 and 4 sequences, respectively, followed by next-generation amplicon sequencing.

### Quantitative PCR (qPCR) analysis

Primers for qPCR analysis (Table S1) were designed using Primer3Plus (https://www.primer3plus.com/index.html). Primer binding efficiencies and specificities were determined using melting curve analysis. To generate cDNA, total RNA was extracted from 4 days post-fertilization (dpf) whole larvae, previously genotyped from tail clips, using Trizol Reagent (Thermo Fisher Scientific) and the RNA Clean and Concentrator Kit (Zymo Research). SuperScript III or IV (Thermo Fisher Scientific) was used to synthesize cDNA from 1 μg of total RNA. cDNA was quantified on a Nanodrop (Thermo Fisher) and diluted to 100 ng/μl. Quantitative PCR was performed using Fast SYBR® Green PCR Mastermix (Thermo Fisher Scientific) on a Quant Studio3 Real-Time PCR System (Thermo Fisher Scientific) using 100ng cDNA per reaction. The ΔΔCt method^38^ was used to calculate fold changes using *rps11* as a reference gene. At least three biological replicates, including three technical replicates each, were included for every experimental group.

### Colorimetric whole-mount in situ hybridization

Whole-mount in situ hybridization was performed in glass vials as described^39^. Riboprobe templates encompassing the first kilobase of coding sequence for *naa15a* and *naa15b* were PCR amplified (Table S1) from a 24-hour post-fertilization (hpf) whole-embryo cDNA library and TOPO-cloned into pCR4 (Thermo Fisher Scientific). Digoxigenin (DIG) probes were synthesized with the DIG RNA labeling kit (Sigma-Aldrich). Anti-sense riboprobes were generated by linearizing templates with *PmeI* and transcribing with T7 polymerase. Sense riboprobes were generated by linearizing the same templates with *NotI* and transcribing with T3 polymerase. Anti-DIG-AP and NBT/BCIP (Sigma-Aldrich) were used to visualize the hybridization signals. Embryos were mounted in 3% methylcellulose in a depression slide prior to imaging under a dissecting microscope.

### Whole-mount immunofluorescence

Whole-mount immunofluorescence was performed as described^40^. The following antibodies were used: Rabbit anti-RCFP (1:200, Takara Bio) to detect ZsYellow, Chicken anti-GFP (1:1000, Aves Labs), Mouse anti-Alcama [(1:50, ZN-8, Developmental Studies Hybridoma Bank (DHSB)], Mouse anti-striated-muscle Tropomyosin (1:200, CH1, DHSB), Rabbit anti-Elastin b (Elnb, 1:1000, formerly Elastin 2)^41^, Mouse anti-sarcomeric Myosin heavy chain (MHC, 1:50, MF20, DSHB), and Mouse anti-cardiac Troponin T (cTNT, 1:1000, CT3, DHSB). The following Thermo Fisher Scientific secondary antibodies were used at 1:500 dilutions: Goat anti-Rabbit IgG 488 to detect ZsYellow and Elastin b, Goat anti-Chicken IgY 488 to detect GFP, Goat anti-Mouse IgG1 647 to detect Alcama and Tropomyosin, Goat anti-Mouse IgG2b 546 to detect MF20, and Goat anti-Mouse IgG1 555 to detect Troponin. Animals were mounted in 0.9% low-melting-point agarose in 35 mm glass-bottom petri dishes (Mattek) prior to confocal imaging.

### Functional analysis of larval hearts

Larvae carrying the *myl7:nucGFP* (OE^myo^ experiments) or *nkx2.5:ZsYellow* (all other experiments) transgenes on 4 days post-fertilization (dpf) were anesthetized with low-dose Tricaine (0.16%, Sigma Aldrich) and mounted on their backs in 0.9% low-melt agarose on a depression slide. Fluorescent hearts were imaged for 15 seconds under a dissecting microscope. The areas (A) of each cardiac chamber were determined during end-diastole (A^ED^) and end-systole (A^ES^) by manually tracing the outer chamber walls during each end-phase and measuring the enclosed perimeters with Fiji^42^. Differences in absolute chamber areas between experiments reflect differences in magnifications used, which were consistent within each experiment. Each data point is the average of measurements from at least 4 cardiac cycles per video. Percent fractional area changes (%FACs) were calculated with the formula 100*(A^ED^-A^ES^/A^ED^).

### Cardiomyocyte quantification and EdU incorporation

Cardiomyocyte (CM) nuclei, with and without replicated DNA, were identified using the *myl7nucGFP* transgene and the Click-iT Plus EdU Imaging Kit (Thermo Fisher Scientific) with adaptations to the manufacturer’s protocol. Larvae were exposed between 3 dpf (72 hpf) and 4 dpf (96 hpf) to 1mM 5-Ethynyl-2’-deoxyuridine (EdU) by adding EdU to the E3^28^ in petri dishes and incubating at 28.5°C. Animals were washed 3 times (3X) with room temperature (RT) E3 and fixed overnight (O/N) at 4°C in 4% paraformaldehyde (PFA). Larvae were washed 3X for 10 minutes (min) in RT PBST [(0.1% Tween-20 in phosphate-buffered saline (PBS), pH=7.4)] before permeabilizing with PBSTx (1% Triton in PBS) for 2 hours (h) at RT. The permeabilization and subsequent steps were performed with agitation. After permeabilization, larvae were washed in RT PBST 3X for 10 min and incubated in Click-iT Plus reaction cocktail for 1h at RT in the dark. Animals were processed to visualize GFP by immunohistochemistry, as described^40^, and imaged by confocal microscopy. Total CM nuclei and CM proliferation indices [(EdU+ CM nuclei/total CM nuclei)*100] in the ventricular and atrial chambers were quantified in confocal Z-stacks using the manual cell counter tool in Fiji^42^.

### Whole-mount TUNEL staining

TUNEL staining was performed with the *In Situ* Cell Death Detection Kit (Fluorescein; Millipore Sigma) with adaptations to the manufacturer’s protocol. Larvae on 4 dpf were fixed O/N at 4°C in 4% PFA, washed 5X in RT PBST for 10 min each, dehydrated in a series of RT methanol washes (25%, 50%, 75% diluted in PBST, and 100%) for 5 min each, and stored O/N at -20°C. Animals were rehydrated in a series of RT methanol washes (75%, 50%, and 25% diluted in PBST) for 5 min each and washed 3X in RT PBST for 5 min each. Larvae were permeabilized with 20 μg/mL Proteinase K for 30 min at RT and washed 3X in PBST for 5 min each. Animals were incubated in prechilled ethanol:acetic acid (2:1) for 10 min at - 20°C and washed 3X in RT PBST for min each. Larvae were incubated in 100 μl TdT Enzyme/Label solution (90 μl Reaction Buffer with 10 μl Tdt Enzyme Solution) O/N at 37°C. Animals were washed for several h in RT PBST, changing the solution every 20 min. Larvae were then processed for immunohistochemistry to detect GFP, essentially as described, before mounting them in 0.9% low-melting-point agarose in 35 mm glass-bottom petri dishes (Mattek) and imaging with confocal microscopy.

### Image analysis of ventricular CM areas and OFT diameters

CM size was measured in the outer and inner curvatures of 4 dpf animals stained for Alcama by manually tracing cell perimeters and measuring the enclosed area with Fiji^42^. For all cells, the optical section showing the maximal cell area was identified and measured. OFT diameter measurements were made in confocal Z-stacks as described^43^. Each data point is the average of 5 serial measurements of the widest OFT diameter.

### Acid-free alcian blue staining

Acid-free alcian blue staining was performed as described^44^ with minor modifications. Four dpf zebrafish larvae were fixed in 4% PFA O/N at 4°C. Animals were washed 3X in PBST (0.1% Tween-20 in PBS) and once with 50% ethanol/PBST for 10 min each at RT with agitation. Larvae were incubated in pre-filtered Alcian Blue staining solution (0.02% Alcian blue, 190 mM MgCl_2_, 70% ethanol) O/N at RT with agitation. Animals were washed 3X in water for 5 min each at RT with agitation and incubated in bleach solution (0.8% KOH, 0.9% H_2_O_2_, and 0.1% Tween-20) for 90 min or until all pigmentation was removed. Larvae were fixed again in 4% PFA at 4°C for 15 min, rinsed in water, and transferred to glycerol. Animals were mounted in glycerol in a depression slide prior to brightfield imaging under a dissecting microscope.

### RNAScope whole-mount in situ hybridization

Whole-mount RNAScope in situ hybridization was performed essentially as described^45^. Briefly, 4 dpf WT *Tg(myl7:nucGFP)* larvae were fixed in 4% PFA for 30 min. Animals were dehydrated in a series of methanol washes (25%, 50%, 75%, 100%) before air drying. Larvae were permeabilized with Pretreat 3^45^ for 20 min with agitation. Whole-mount fluorescence in situ hybridization was performed using the RNAscope Multiplex Fluorescent Reagent Kit v2 (ACDBio), followed by whole-mount immunofluorescence to detect GFP and counterstaining with DAPI. Larvae were mounted in 0.9% low-melting-point agarose in 35mm glass-bottom Petri dishes (Mattek) and imaged with a confocal microscope. The riboprobes targeted proprietary 20 basepair (bp) sequences within bp 202-1429 of *naa15a* (NM_200646.2) or bp 304-1366 of *naa15b* (NM_203321.2).

### Generation of *myl7:nuc-mCherry-P2A-HA-naa15a* transgenic zebrafish

The *myl7:nuc-mCherry-P2A-HA-naa15a* transgene was generated by HiFi DNA Assembly Cloning Kit (New England Biolabs). A cDNA encoding full-length Naa15a, HA tagged on the N-terminus, was ordered as a gBlock^TM^ (IDT DNA) and cloned in frame with nuclear(nuc)-mCherry-P2A-encoding sequences, all downstream of the ∼0.9 kb myocardial *myl7* promoter^46^ in a Tol2-based backbone. Additional cloning details and plasmid sequence available upon request. Fifty picograms (pg) of the transgene were co-injected with 100 pg of Tol2 mRNA into the cells of one-cell stage embryos. Germline transmission was monitored by F1 heart fluorescence. The resulting OE^myo^ transgenic strain and WT control animals, both carrying the *myl7:nucGFP* transgene, were assessed on 4 dpf for %FACs as described above.

Human *NAA15* mRNA synthesis, microinjections, and variant testing.

For in vitro transcription, a cDNA encoding human NAA15 (Ensembl Gene ID: ENSG00000164134; NM_057175) with a C-terminal HA tag was PCR amplified (Table S1) from pCMV6-*NAA15* (OriGene, RC210884). The PCR product was cloned into pCS2+^47^ using the HiFi DNA Assembly Kit (New England Biolabs) after digestion of the vector backbone with *ClaI* and *XhoI*. The cDNAs encoding protein variants K336fs, R267W, and L566V^6^ were generated by site-directed mutagenesis of pCMV6-*NAA15* using the Q5 Site-Directed Mutagenesis Kit (New England Biolabs; Table S1) according to the manufacturer’s instructions, sequence verified (Table S1), and subcloned into pCS2+, as above. The mCherry cDNA was PCR amplified (Table S1) and cloned into pCS2+ using the Quick Ligation Kit (New England Biolabs) after digesting the amplicon and vector backbone with C*laI* and *XhoI*. For in vitro transcription, pCS2+*-mCherry* was linearized with *SacII*. The pCS2+ plasmids with WT and variant *NAA15* cDNAs were linearized with *NotI*. All mRNAs were transcribed with the SP6 mMessage mMachine Kit (Thermo Fisher Scientific). The resulting mRNAs were purified using the RNA Clean & Concentrator™ Kit (Zymo Research), and 100 picograms of each were injected into the yolks of one-cell stage embryos. Percent FACs were measured on 4 dpf as described above.

### Analysis of *naa15^RD^* adult animals, hearts, and CMs

Following echocardiography analysis (See below), WT and *naa15*^RD^ animals at 9 months post-fertilization (mpf) were euthanized by ice bath immersion, sized by weight and length, and their dissected hearts were imaged under a dissecting microscope in a standardized orientation before fixation O/N at 4oC in 4% PFA. Ventricular size was determined by manually tracing the ventricular perimeter in photographs and measuring the enclosed area with Fiji^42^. Histological sectioning and AFOG staining were performed as described^31^. Single-cell spreads of CMs from 9 mpf WT and *naa15RD* hearts were generated and dried as described^31^. Spreads were rehydrated in PBS, permeabilized/blocked with 0.1% IGEPAL/3% BSA/PBS for 45 min at RT, and incubated O/N at 4°C in primary antibody [mouse anti-striated-muscle Tropomyosin (1:500, CH1, DHSB) in 3% BSA/PBS)]. Slides were washed 3X in RT PBS for 5 min each and incubated for 1h at RT in secondary antibody [goat anti-Mouse IgG1 568 (Thermo Fisher Scientific) diluted 1:500 in 3% BSA/PBS]. Slides were washed 3X in RT PBS for 5 min each, counterstained in DAPI, mounted in FluorSave reagent (Calbiochem) and imaged using a confocal microscope. CM size was determined by manually tracing the cell perimeter and measuring the enclosed area in Fiji^42^. Cells were also categorized as “organized” or “disorganized” based on qualitative assessments of myofibrillar/sarcomere structure.

### Echocardiography

Images were acquired in B-mode, color Doppler, and pulse wave Doppler modes using the Vevo 3100 ultrasound system with a MX700 ultrasound transducer at a frequency of 40 mHz as previously described^48–50^. Zebrafish were lightly anesthetized in 0.4 mg/mL Tricane for 45 seconds, suspended ventral side up in a slit sponge, and submerged in 0.4 mg/mL Tricane for the duration of imaging, approximately 2 – 5 min. The ultrasound transducer was positioned parallel to the A-P axis of the zebrafish at an approximately 70° angle. Peak ventricular outflow (V_o_^PK^) regions were located during the live imaging session in color Doppler mode. Outflow measurements were then detected, with angle correction, using pulse wave Doppler at the bulboventricular valve. Heart rate and peak ventricular outflow velocities were manually calculated in the Vevo analysis software from the average of at least 3 cardiac cycles. Heart variabilities were determined using the Vevo analysis software by manually measuring the time intervals between 6 successive heartbeats (i.e., interbeat intervals identified by V_o_^PK^) and calculating a standard deviation for each heart. To determine ventricular areas during end-systole (VAs) and end-diastole (VAd) as well as end-diastolic (EDV) and end-systolic (ESV) volumes, B-mode images of the ventricular wall were manually outlined in Vevo using the area and LV trace tools. Ejection fraction (EF) was calculated as EF = (EVD – ESV)/EDV. Stroke volume (SV) was determined by SV = EDV-ESV. Cardiac output (CO) was calculated as CO = HR x SV, where HR is heart rate.

### Sample processing for proteomics analysis

Ventricles from 9 mpf WT and *naa15^RD^* animals were dissected and pooled into 5 replicates per cohort, 3-5 ventricles per replicate. Ventricles were suspended in cold, freshly-prepared lysis buffer consisting of 5% Sodium Dodecyl Sulfate (ThermoFisher), 50 mM triethylammonium bicarbonate buffer (Sigma-Aldrich), 50 mM magnesium chloride (Sigma-Aldrich), containing protease inhibitors, and incubated for 5 min on the benchtop. Samples were then centrifuged for 10 min at 4°C and 18000 rcf to remove debris. Supernatant was removed, and protein concentrations were determined by BCA assay (ThermoFisher). Proteins were reduced with 5 mM dithiothreitol (DTT, Sigma-Aldrich), followed by alkylation with 10 mM iodoacetamide (IAA, Sigma-Aldrich). Samples were then acidified with phosphoric acid for a final concentration of 1.2% and loaded onto midi S-traps (Protifi) in a ratio of 1:6 with chilled S-trap buffer (90% methanol, 100 mM triethylammonium bicarbonate buffer). Proteins were digested with digestion buffer containing 1:40 enzyme:substrate ratio of LysC endopeptidase (Wako Chemicals) and sequencing grade modified trypsin (Promega) O/N. Samples from S-trap cartridges were then eluted, dried using a vacuum concentrator, and desalted using Sep-Pac C18 SPE cartridges (Waters). Clean peptides were dried using a vacuum concentrator, and peptide concentrations were determined by BCA assay. 100 μg aliquots of every sample were prepared and dried down for subsequent labeling using Tandem Mass Tag (TMT) 16-plex reagent. Samples were randomized into 15 of the TMT 16-plex channels and resuspended in 20 μL of 50 mM HEPES pH 8.5. 20μL of each TMT reagent was added to the corresponding sample. Labeling was allowed to proceed for 1 h in a thermomixer set to 25°C and 1000 rpm. After 1 h, a small aliquot was removed from each sample for labeling efficiency and mixing tests. Following QC checks, reactions were quenched with 5% hydroxylamine, and samples were combined, dried using a vacuum concentrator, and desalted using Sep-Pac C18 SPE cartridges (Waters). Desalted and dried down sample was reconstituted in 5 mM ammonium formate, pH 10, and fractionated by high pH reversed phase chromatography using a 3.5 μm Agilent Zorbax 300 Extend-C18 column (4.6 mm ID x 250 mm length). Sample was eluted over 96 min gradient, collecting fractions every minute. Fractions were then concatenated into 24 total fractions and analyzed by LC-MS/MS.

### LC-MS/MS analysis of samples

Samples were analyzed on an Orbitrap Exploris 480 mass spectrometer (Thermo Fisher Scientific) coupled with a Vanquish Neo UHPLC system (Thermo Fisher Scientific). An in-house packed 26 cm x 75 μm internal diameter C18 silica picofrit capillary column (New Objective) with 1.9 mm ReproSil-Pur C18-AQ beads (Dr. Maisch GmbH, r119.AQ) heated at 50°C was used for online LC-MS/MS. Samples were separated with a Solvent A composed of 0.1% formic acid (FA) in water and Solvent B composed of 0.1% FA in acetonitrile (ACN) at the flow rate of 200nL/min. One microgram of each fraction was loaded on-column and eluted with a 110-min LC-MS/MS method consisted of the following gradient profile:1.8-5.4% B in 1 min, 5.4-27% B in 84 min, 27-54% B in 10 min, 54-81% B in 1 min, followed by a hold at 81% B for 5 min, drop to 45% B, and a hold at 45% B for 9 min. Data-dependent MS/MS acquisition was performed using the following parameters: positive ion mode at a spray voltage of 1.8 kV; MS1 resolution of 60,000, normalized AGC target of 300%, a mass range from 350 to 1800 *m/z*, and maximum injection time of 10ms; top 20 ions were selected with isolation window of 0.7 *m/z,* MS2 resolution of 45,000, normalized AGC target of 30%, and maximum injection time of 105 ms; HCD collision energy of 32%; dynamic exclusion for 15 seconds, and charge states 2-6. Precursor fit filter of 50% was used with a fit window of 1.2 m/z.

### Data Analysis

The raw MS/MS data was searched on Spectrum Mill MS Proteomics Software (Broad Institute) using UniProt zebrafish database downloaded on 11/04/2020, with digestion enzyme conditions set to “Trypsin allow P”, < 4 missed cleavages, cysteine carbamidomethylation and TMT16 on N-term and lysine as fixed modifications and oxidized methionine, acetylation of the protein N-terminus, pyroglutamic acid on N-term Q, and pyro carbamidomethyl on N-term C, deamidation of N as variable modifications. Matching criteria included a 40% minimum matched peak intensity and a precursor and product mass tolerance of +/- 20 ppm. Peptide-level matches were validated if found to be below the 1.2% false discovery rate (FDR) threshold and within a precursor charge range of 2-6. A second round of validation was then performed for protein-level matches for proteome datasets, requiring a minimum protein score of 13 and protein level FDR of 0%. Protein quantification was achieved by taking the ratio of TMT reporter ions for each sample over the median of all channels. TMT16 reporter ion intensities were corrected for isotopic impurities in the Spectrum Mill protein/peptide summary module using the afRICA correction method, which implements determinant calculations according to Cramer’s Rule79 and correction factors obtained from the reagent manufacturer’s certificate of analysis for lot number VE299607. After performing Median-MAD normalization of the proteome dataset, a moderated two-sample t-test was applied to the dataset to compare sample groups. Benjamini-Hochberg corrected p-value thresholds were used to assess differentially expressed proteins between experimental conditions.

For N-terminome analysis, peptide-level results exported from Spectrum Mill were filtered to retain peptides with N-terminal start positions of 1 or 2, a deltaForwtoRev > 0, and a peptide score > 7. Peptides containing missing values were excluded. The resulting dataset comprised 742 peptides, the majority of which were N-terminally acetylated, and was used for quantitative analysis of N-terminal acetylation. TMT16 reporter ion ratio calculation and statistical analyses were performed as described above for the global proteome.

### Gene Ontology term enrichment analysis

To perform Gene Ontology analysis, the UniProt IDs were mapped to gene symbols using annotations downloaded from the UniProt website on 6/2/25. The obtained gene symbols for up-and down-regulated proteins were used for GO enrichment analysis, respectively, using the enrichGO function in the clusterProfiler package (version 4.0.5) in R. All detected protein names were used as background. The up- and down-regulated proteins were defined as those with adjusted p-values < 0.05 and |Fold Change (FC)| > 1. The total number of genes in the GO term ‘mitochondrial respiratory chain complex’ was obtained online from AmiGO 2 (http://amigo.geneontology.org/; version 2.5.17; data accessed in February 2026)^51^.

### Cross-species comparisons of differentially expressed proteins downstream of *NAA15*

UniProt accession numbers for the fully quantified zebrafish proteins (Table 1) were used to identify corresponding gene symbols in the UniProt database^32^, yielding 5031 zebrafish genes. Of these, 4262 had orthologous human genes, based on manually curated orthology data available from ZFIN^52^, downloaded from https://zfin.org/downloads (November 2025).

**Table 1.** Spreadsheets showing the principal component analysis, fully quantified proteins, and differentially expressed proteins (DEPs; adjusted p<0.05, FC>|1|) in *naa15^RD^*hearts as determined by deep quantitative proteomic profiling. The log_2_FC values are reported as both WT/*naa15^RD^* and *naa15^RD^*/WT (gray background).

Proteins identified in human induced pluripotent stem cells, including WT cells and those carrying one (*NAA15^+/-^*) or two (*NAA15^-/-^*) null alleles of *NAA15,* were downloaded from Ward et. al.^6^ Reported p-values for differential protein expression between WT and mutant lines were corrected for multiple testing by calculating false-discovery rates (FDRs), mirroring the correction applied to the zebrafish dataset. Intersecting the 4262 zebrafish gene symbols with the 3812 human gene symbols from Ward et al.^6^ yielded 2013 zebrafish-human orthology pairs (Table 5). Filtering based on adjusted p-value (FDR<0.05) and log2FC enabled the identification of proteins differentially expressed exclusively in zebrafish *naa15^RD^*adult hearts, in human stem cells (*NAA15^+/-^* or *NAA15^-/-^*), or both species, with or without considering the direction of differential expression (UP/DOWN). The Fisher’s exact test was used to determine statistical significance of overlap between datasets. Venn diagrams were generated with ggVennDiagram (version 1.5.4)^53^.

### Transmission electron microscopy

Larvae at 4 dpf were fixed in 2.5% glutaraldehyde, 1.25% PFA, 0.03% picric acid in 0.1M sodium cacodylate buffer (pH=7.4) O/N at 4°C. Fixed animals were washed in 0.1M sodium cacodylate buffer, postfixed with 1% osmium tetroxide/1.5% potassium ferrocyanide for 1 h, washed 2X in water, 1X in Maleate buffer (MB), incubated in 1% uranyl acetate in MB for 1 h, and washed 2X in water. Larve were dehydrated in a series of methanol washes (50%, 70%, 90%, 2X 100% methanol] for 10 min each. The samples were incubated in propylene oxide for 1 h and infiltrated O/N in a 1:1 mixture of propylene oxide and TAAB (TAAB Laboratories Equipment Ltd). The following day, samples were embedded in TAAB Epon and polymerized for 48 h at 60°C. Ultrathin sections of approximately 60nm were cut on a Reichert Ultracut-S microtome, transferred to copper grids, stained with lead citrate, and examined under a Tecnai G^2^ Spirit BioTWIN or JEOL 1200EX transmission electron microscope. Images were captured with an AMT 2k CCD camera.

### Fluorescence Imaging of Isolated Hearts

Hearts were isolated from 4 dpf zebrafish larvae and placed in normal Tyrode’s solution (NTS) containing 136□mM Na⁺, 5.4□mM K⁺, 1.0□mM Mg²⁺, 0.3□mM PO₄³⁻, 1.8□mM Ca²⁺, 5.0□mM glucose and 10.0□mM HEPES at pH□7.4, supplemented with 8 mg/ml BSA. To assess relative mitochondrial membrane potential (ΔΨ*_m_*), hearts were loaded with 2.5 μM of the ratiometric dye JC-1 for 15 minutes, followed by a 15-minute washout in NTS.

Prepared hearts were transferred to a chamber (RC-49MFS; Warner Instruments) perfused with NTS supplemented with 1 mmol/L cytochalasin D (Sigma) to decouple electrical impulses from mechanical contractions. The chamber was mounted onto the stage of an inverted microscope (TW-2000; Nikon) equipped with wires for cardiac pacing. Fluorescence was collected by a high-speed 80×80-pixel 14-bit CCD camera (RedShirtImaging). Using a 20× objective and a 0.5× C-mount adapter, the final optical magnification was 10×, resulting in a pixel-to-pixel distance of 2.4 µm. Static two-channel fluorescence images were acquired; JC-1 monomers were visualized in the green channel (470 nm excitation, ∼510-530 nm emission) and highly concentrated J-aggregates were visualized in the red channel (∼530-550 nm excitation, ∼585-590 nm emission).

### Mitochondrial Membrane Potential (ΔΨ*_m_*) Image Analysis

Image processing and ratiometric analyses for JC-1 experiments were performed using a custom-written pipeline in Python. To correct for minor spatial drift between channel acquisitions, red channel images were aligned to the green channel utilizing phase cross-correlation, supplemented by interactive user-defined shifts when necessary. Cardiac tissue was segmented using Otsu thresholding on the intensity of each channel individually and the intersection of these segmentations was then used to generate a single, unified tissue mask. Background fluorescence was estimated independently for both the green and red channels by calculating the 5th percentile of intensity values among non-tissue (background) pixels. This background value was subtracted from the respective full image. A spatial ΔΨ*_m_* map was generated by calculating the pixel-wise red-to-green fluorescence intensity ratio (background-subtracted red / background-subtracted green). ROIs were interactively defined on an aligned composite image and saved for reproducibility. For each ROI, the relative ΔΨ*_m_* was quantified as the average of the per-pixel ratios within the regional boundary.

### Statistics

Prism software (GraphPad) was used to perform statistical tests and generate graphs. Outliers were identified using the ROUT method (Q=1) and removed. Data normality was evaluated using the Shapiro-Wilk test. For normally distributed data, statistical significance was determined by the Student’s t-test for pairwise comparisons or the one-way ANOVA, followed by Tukey’s multiple comparisons tests, when three or more groups were evaluated. For non-normal data, significance was determined by the non-parametric Mann-Whitney Test for pairwise comparisons or the Kruskal-Wallis Test, followed by Dunn’s multiple comparisons test, when three or more groups were evaluated. Differences between experimental groups were considered statistically significant at p<0.05. Additional information, including the specific statistical tests used and sample sizes, is provided in the figure legends.

## Results

### Naa15-deficient zebrafish exhibit diminutive, lowly contractile ventricles and early larval lethality

Zebrafish Naa15a and Naa15b proteins share 84% identity and greater than 80% identity with human NAA15 (Fig. S1A), suggesting that NAA15’s molecular function remains unchanged after 450 million years^54^. Accordingly, the domain structure of NAA15 is highly conserved and characterized by multiple tetratricopeptide repeats, which are typically mediators of protein-protein interactions^55^, and a coiled-coil domain, a common structural motif with numerous functions (Fig. S1B, Fig. S2A)^56^.

Using CRISPR/Cas9-mediated genome editing, we isolated mutant alleles of *naa15a* (*naa15a^chb13^*) and *naa15b (naa15b^chb14^*). The *naa15a^chb13^* allele contains a 12-base pair (bp) deletion (ΔGAACATGGAGGC) encompassing the last 10 bp of exon 2 and a non-canonical splice donor site (GC) in the downstream intron. This deletion would shift the open reading frame if the transcript were correctly spliced. However, given the absence of the splice donor site, we reasoned that the intron might be retained in some percentage of transcripts. This was confirmed by next-generation sequencing of PCR products spanning exons 2 and 3, amplified from double-mutant (*naa15a^chb13/chb13^*; *naa15b^chb14/chb14^*) cDNA. As predicted, 53% of reads (n=14725/27869) were devoid of the 12 bp sequence while retaining the downstream intron. This major mutant mRNA isoform resulted in the replacement of three amino acids (aa) at the start of the first TPR repeat with a 32-aa insertion encoded by the intron (Fig. S2A, Fig. S3A). Additionally, 38% percent of the reads (n=10660/27869) showed no intron retention, but aberrant usage of a cryptic splice donor site (GT) 17 bp upstream of the deleted splice donor site. This second major mutant mRNA isoform is predicted to encode 40 wild-type (WT) and 19 frame-shifted aa, out of 876 total, before a premature stop codon truncates the protein (Fig. S2A, Fig. S3A). The *naa15b^chb14^* allele contains a 5 bp deletion (ΔGGAGA) and a 10 bp insertion (ATTGGACTCC) in exon 7, resulting in a net insertion of 5 bp and a shifted open reading frame. The mutant allele encodes 255 WT and 23 frame-shifted aa, out of 865 total, before a premature stop codon truncates the protein (Fig. S2A, S3B).

We performed quantitative PCR (qPCR) analysis on 4 days post-fertilization (dpf) single-and double-mutant animals to determine how the indels affected mRNA stability, predicting that the mutant mRNAs with premature stop codons could be susceptible to nonsense-mediated decay^57^. The retained intron’s effect on message stability was harder to predict because intron retention can have variable effects on transcript stability^58,59^. Single mutants showed reduced levels of the mutant transcripts, without compensatory upregulation of the paralogous genes (Fig. S2B). In double mutants, both transcripts were severely reduced, approaching log_2_-fold changes of 0 (Fig. S2B). Importantly, the primers to detect *naa15a* amplified across an exon-exon junction downstream of the retained intron to ensure that its retention didn’t interfere with transcript detection. The significantly reduced stability of both *naa15* transcripts in double mutants strongly suggests that they are *naa15* double knockout (DKO) animals.

We performed qPCR analysis on DKO animals for genes encoding other members of Naa15-containing complexes^16^. We detected upregulation of genes encoding the catalytic subunit of NatA (Naa10) and the NatA-interacting protein Hypk, suggesting that DKO animals attempted to compensate for severely reduced or absent NatA activity (Fig. S2C). Expression of Naa50, the catalytic subunit of NatE complex, of which Naa15 is also a member, was unchanged in DKO animals (Fig. S2C), suggesting that NatE activity was less affected by deletion of *naa15*.

Single-mutant *naa15a^chb13/chb13^* and *naa15b^chb14/chb14^* animals were adult-viable, fertile, and devoid of gross phenotypes. By contrast, *naa15a^chb13/chb13^*; *naa15b^chb14/chb14^* DKO larvae displayed several abnormalities, including pericardial edema, a diminutive lower jaw, and no swim bladder (Fig. 1A), all of which appeared by 4 dpf when the maternal dose was minimized (See below). Beyond 4 dpf, the pericardial edema progressed, and DKO animals invariably became necrotic on or before 8 dpf. The observation that these phenotypes emerged exclusively in double mutants reveals the redundant nature of *naa15a* and *naa15b* in zebrafish.

**Figure 1.**
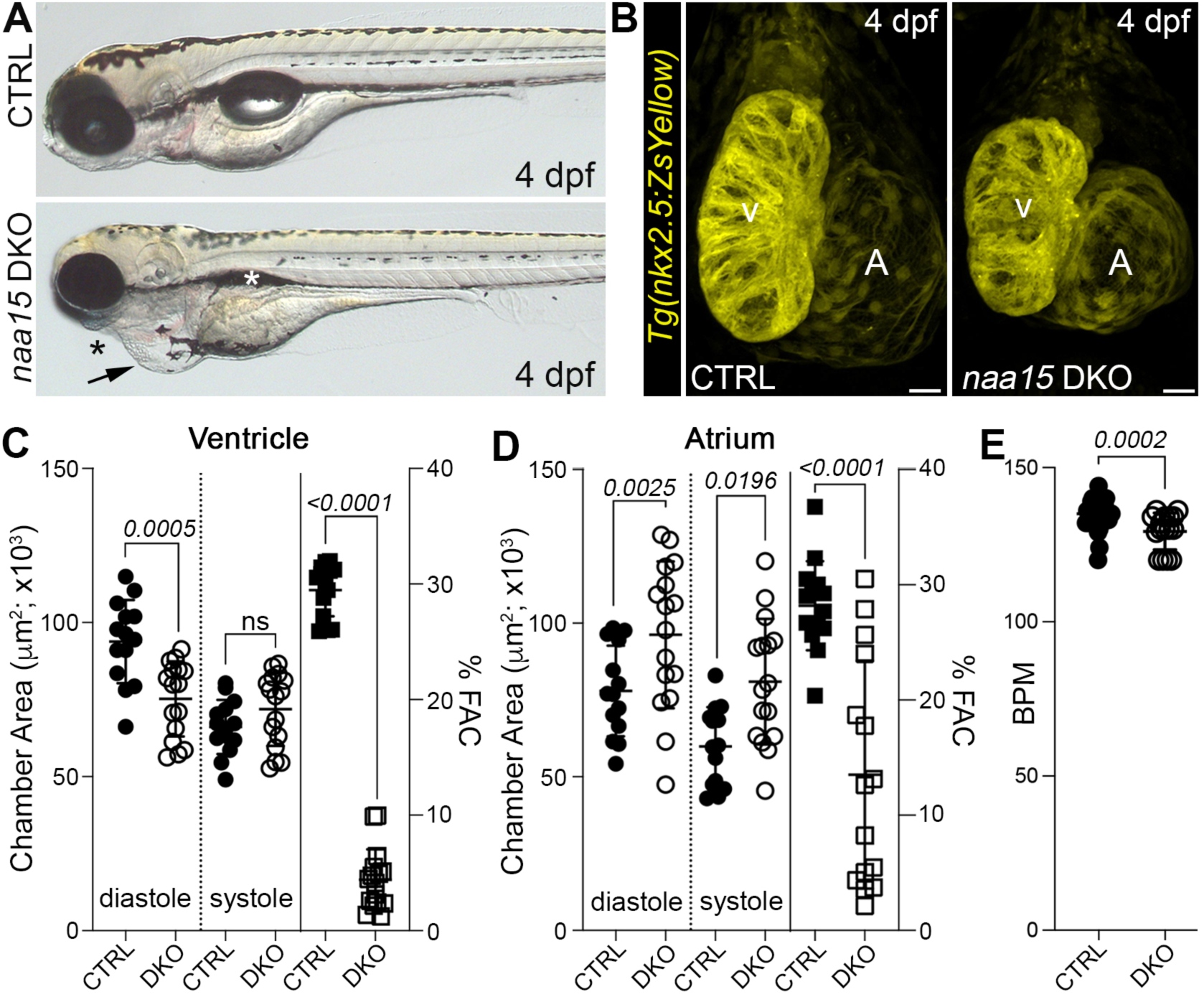
Reduced ventricular size, contractility, and heart rate in *naa15*-deficient zebrafish. (A) Brightfield images of control-sibling (CTRL) and *naa15a; naa15b* double knockout (DKO) zebrafish at 4 days post-fertilization (dpf). The arrow points to pericardial edema, the black asterisk highlights an underdeveloped lower jaw, and the white asterisk indicates an absent swim ladder. (B) Confocal projections of 4 dpf CTRL and DKO hearts in animals carrying the *nkx2.5:ZsYellow* transgene, immunostained for ZsYellow. (C,D) Dot plots showing the ventricular (C) and atrial (D) end-diastolic areas, end-systolic areas, and percent (%) fractional area changes (%FACs) in 4 dpf CTRL (n=14) and DKO (n=16) larvae. (E) Dot plot showing heart rates of CTRL (n=37) and DKO (n=16) animals at 4 dpf. Statistical significance was determined by Student’s t-tests, except for ventricular %FAC and heart rate, which were determined by Mann-Whitney tests. Means ± 1SD are shown. Scale bars=25 μm. Abbr: V, ventricle; A, atrium; ns, not significant; BPM, beats per minute.

During initial phenotyping, we observed that the pericardial edema in DKO animals was variably penetrant, depending on the mother’s genotype, suggesting a maternal effect. Specifically, when the mother carried a single WT allele of either *naa15a* or *naa15b* (i.e. *naa15a^+/-^*; *naa15b^-/-^ or naa15a^-/-^*; *naa15^+/-^*), the pericardial edema was 100% penetrant in DKO offspring at 4 dpf (Fig. S4A; crosses #1-3). By contrast, when the mother was double heterozygous for *naa15a* and *naa15b* (i.e., *naa15a^+/-^*, *naa15b^+/-^*animals), the penetrance was reduced to ∼30% (Fig. S4A; crosses #4,5). The pericardial edema, however, emerged on subsequent days when the penetrance approached 100%. These data suggest that the maternal dose provided by two WT *naa15* alleles, but not one, is sufficient to mask the pericardial edema in ∼70% of 4 dpf DKO offspring. We confirmed by in situ hybridization that both transcripts are maternally deposited (Fig. S4B). To ensure consistent timing of phenotypic emergence, we analyzed larvae exclusively from mothers carrying a single WT copy of either *naa15a* or *naa15b*.

Because pericardial edema is often a sensitive indicator of cardiac defects^60^, we examined hearts in 4 dpf control-sibling (CTRL) and DKO animals carrying the *nkx2.5:ZsYellow* transgene^30^, which labels chamber myocardium at this stage^61^. In fixed larvae immunostained for ZsYellow, DKO ventricles were smaller than those in CTRL animals, whereas atrial size appeared largely unaffected (Fig. 1B). To measure cardiac contractility, we captured video images of CTRL and DKO hearts, measured chamber areas during end-systole and end-diastole, and calculated percent (%) fractional area changes (FACs). Compared to CTRL ventricles, DKO ventricles were smaller during diastole, contracted minimally during systole, and exhibited an 85% decrease in %FAC (Fig. 1C). By contrast, the atria in live DKO animals were dilated, likely a secondary consequence of reduced ventricular function^62^, but also displayed a reduction in %FAC (Fig. 1D). Lastly, DKO hearts showed a modest but statistically significant reduction in heart rate (Fig. 1E). These data demonstrate that *naa15* is essential for establishing or maintaining proper cardiac chamber size, contractility, and heart rate in larval zebrafish.

### naa15 is required for ventricular cardiomyocyte proliferation, growth, and myofibril optimization

To determine if the chamber-size abnormalities in DKO animals resulted from deviations in cardiomyocyte (CM) number or size, we first quantified CMs in 4 dpf CTRL and DKO animals carrying the *myl7:nucGFP* transgene, which expresses GFP in CM nuclei^31^. DKO ventricles contained 50% fewer CMs than WT ventricles (Fig. 2A,B). Mutant atria were also hypocellular (Fig. 2A). To learn if these deficits stemmed from decreased CM cell division, we exposed CTRL and DKO *Tg(myl7:nucGFP)* animals to the thymidine analog 5-Ethynyl-2’-deoxyuridine (EdU) between 3 and 4 dpf, quantified total and EdU+ CM nuclei at 4 dpf, and calculated proliferation indices for both chambers. Whereas the index exceeded 5% in the ventricles and atria of CTRL animals, DKO CMs showed little to no cell cycle activity (Fig. 2C,D), demonstrating that *naa15* is indispensable for CM proliferation. Using TUNEL staining, we ruled out CM apoptosis as a contributing factor to CM hypoplasia (Fig. S5A). To quantify CM size, we measured ventricular CM areas in optical sections of 4 dpf CTRL and DKO ventricles after immunostaining with the cell-surface marker Alcama^36^. Ventricular CMs in DKO animals were significantly smaller by approximately 75% in both the inner and outer curvatures (Fig. 2E,F), which also contributed to the reduced ventricular size.

**Figure 2.**
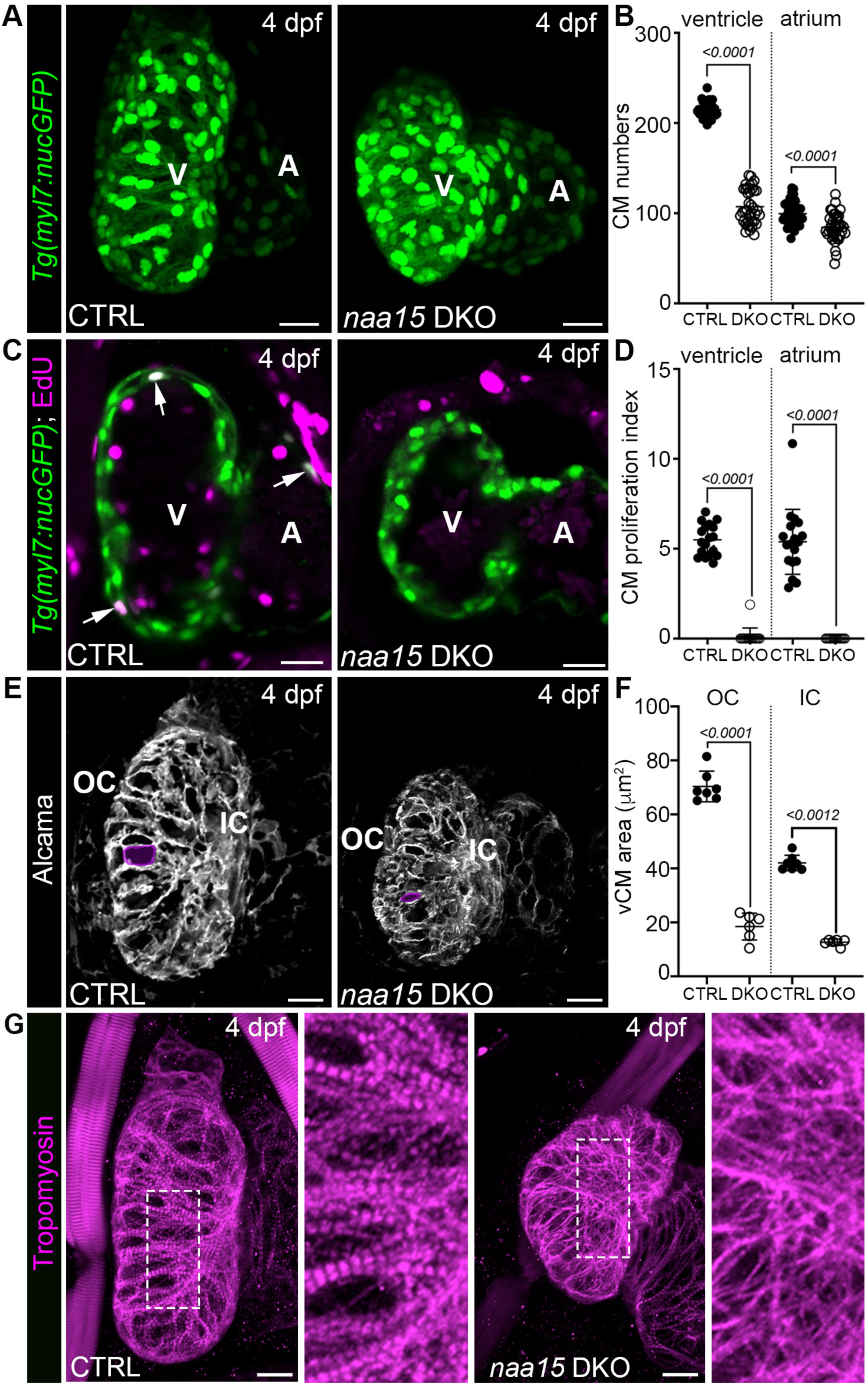
Reduced CM number, proliferation, cell size, and myofibril organization in *naa15*-deficient zebrafish. (A) Confocal projections of hearts in 4 days post-fertilization (dpf) control-sibling (CTRL) and double knockout (DKO) zebrafish carrying the *myl7:nucGFP* transgene, immunostained for GFP. (B) Dot plot showing ventricular and atrial cardiomyocyte (CM) numbers in CTRL (n=40) and DKO (n=38) animals at 4 dpf. (C) Single optical sections of hearts in 4 dpf CTRL and DKO animals carrying the *myl7:nucGFP* transgene, exposed to EdU between 3 dpf and 4 dpf, and double immunostained for GFP and EdU. Arrows highlight EdU+ CM nuclei in the ventricle and atrium of the CTRL heart. (D) Dot plot showing the ventricular and atrial CM proliferation indices in CTRL (n=18) and DKO (n=16) animals at 4 dpf. (E) Confocal projections of hearts in 4 dpf CTRL and DKO animals, immunostained for the cell surface marker Alcama. Single CMs are outlined as representative examples of cellular areas. (F) Dot plot showing ventricular CM (vCM) areas in the outer curvatures (OCs) and inner curvatures (ICs) of CTRL (n=7) and DKO (n=6) hearts at 4 dpf. Each data point is the average of six cell areas per region from each heart. (G) Confocal projections of CTRL and DKO hearts in animals immunostained for the thin filament protein Tropomyosin. All CTRL hearts (n = 8/8) exhibited lateral bundling of and longitudinal alignment of myofibrils, creating prominent striations. For DKO hearts, approximately half (n=8/14) were indistinguishable from WT (not shown), but the rest (6/14) showed moderate myofibril disarray and reduced striations. For all dot plots, statistical significance was determined by Mann-Whitney tests, except for atrial cell numbers and OC vCM areas, which were determined by Student’s t-tests. Means ± 1SD are shown. Scale bars=25 μm. Abbr: V, ventricle; A, Atrium.

Next, we visualized cardiac myofibrils and their sarcomeres by immunostaining 4 dpf CRTL and DKO animals with an antibody against the thin-filament protein tropomyosin^63^. In WT ventricles, bundles of aligned myofibrils with prominent striations were observed in a mesh- or sponge-like pattern across the ventricle (Fig. 2G). In DKO animals, myofibrils were present, but in ∼50% of animals, they showed reduced consolidation into discrete bundles, and their striations were less prominent based on qualitative assessments (Fig. 2G). Taken together, these data demonstrate that *naa15* is required for ventricular growth by promoting CM cellular growth and proliferation, while also promoting or fine-tuning myofibril organization.

Lastly, we investigated DKO animals for outflow tract (OFT) and pharyngeal arch artery (PAA) phenotypes, which are known to coexist with ventricular defects in both zebrafish and higher vertebrates, due to their shared embryonic origins and dependencies on key developmental pathways^39,64–69^. On 4 dpf, DKO OFTs were indistinguishable from those in CTRL animals based on general morphology, smooth muscle marker expression (i.e., Elnb), and cross-sectional diameter (Fig. S5B,C). Similarly, PAAs III-VI were unaffected by loss of *naa15* (Fig. S5D). These data demonstrate that among several structures that derive from a previously characterized *nkx2.5+* progenitor pool in zebrafish^39,61,65,70,71^, the ventricle is uniquely affected by *naa15* deficiency, suggesting that *naa15* functions in ventricular CMs (See below) rather than a progenitor population.

### Naa15 mutants display craniofacial abnormalities involving pharyngeal cartilage and muscle

Given that DKO animals show gross underdevelopment of the lower jaw (Fig. 1A), we evaluated the pharyngeal cartilage and muscles in CTRL and DKO animals at 4 dpf. Alcian blue staining revealed the presence of bilateral cartilage segments in pharyngeal arches (PA) I and II, but they were undersized and displaced posteriorly. This was readily apparent by comparing their positions relative to the ethmoid plate and eyes in both groups (Fig. 3A). In the posterior PAs, the bilateral cartilage segments were similarly displaced (Fig. 3A). Examination of the pharyngeal muscles revealed that the ventral and middle PAI- and PAII-derived muscles^65^ were present, but significantly shorter in the anterior-posterior axis, with the PAI-derived muscles being most severely affected. (Fig. 3B). These data demonstrate that *naa15* is required for the growth or anterior-posterior positioning of pharyngeal cartilage and muscle, consistent with the presence of mild craniofacial dysmorphologies in some patients with *NAA15*-related syndrome^9,13^.

**Figure 3.**
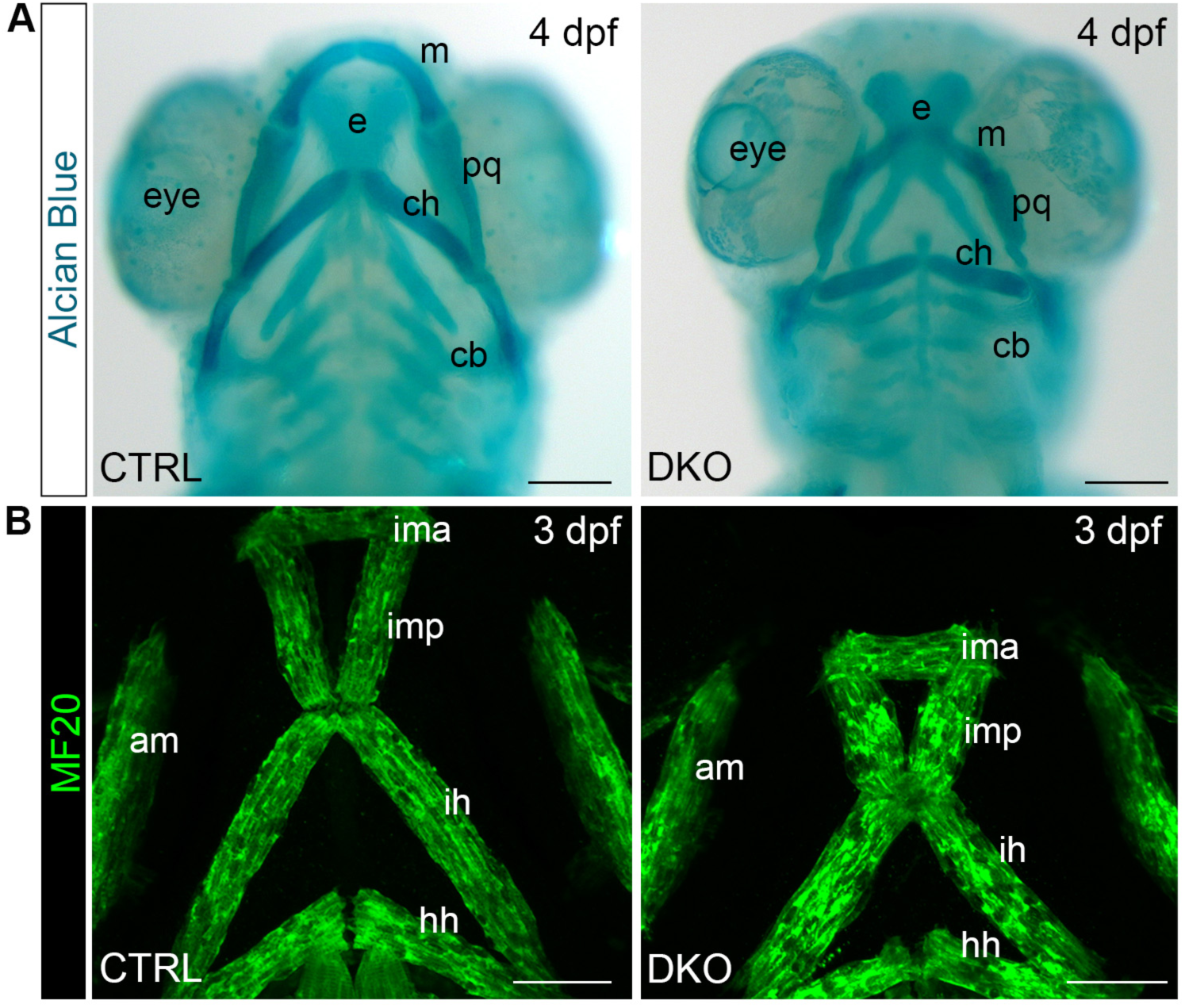
Defects in the growth or positioning of pharyngeal cartilage and muscles in *naa15*-deficient zebrafish. (A) Brightfield ventral views of 4 days post-fertilization (dpf) control-sibling (CTRL) and double knockout (DKO) animals stained with Alcian Blue to visualize cartilage. (B) Confocal projections of ventral head muscles in 3 dpf CTRL (n=5) and DKO (n=6) animals stained for striated muscle (MF20 antibody). For both (A) and (B), similar staining patterns were observed across all animals in each experimental group. Abbr: V, ventricle; A, atrium; OFT, outflow tract; ventral PA1 (mandibular) muscle: ima, intermandibular anterior; middle PAI muscles: imp, intermandibular posterior and am, adductor mandibulae; ventral PAII (hyoid) muscle: ih, interhyal; middle PA2 muscle: hh, hyohyal; VA, ventral aorta; PAI cartilage: m, meckel’s and pq, palatoquadrate; PAII cartilage: ch, ceratohyal; PAIII-VII cartilage: cb, ceratobranchial cartilage.

### Myocardial Naa15 re-expression partially rescues ventricular contractility and growth

The developmental expression patterns of *naa15a* and *naa15b* in 24 hours post-fertilization (hpf) zebrafish embryos were previously reported to include the eye, midbrain, and somites^19^. Nonetheless, an analysis of later-stage expression, which is relevant to the DKO heart phenotype, has not been reported. Therefore, we performed RNAScope in situ hybridization for both genes on 4 dpf animals expressing GFP in CM nuclei. Examination of single optical sections revealed localization of both transcripts to the ventricular myocardium, atrial CMs, and adjacent OFTs (Fig. 4A,B). When DKO animals were processed in parallel, signals were virtually absent (Fig. 4B), simultaneously corroborating riboprobe specificities and our qPCR analysis (Fig. S2B). These data demonstrate that *naa15* genes localize prominently to the ventricular myocardium of early larval zebrafish at a stage when DKO animals exhibited ventricular abnormalities (Fig. 1; Fig. 2).

**Figure 4.**
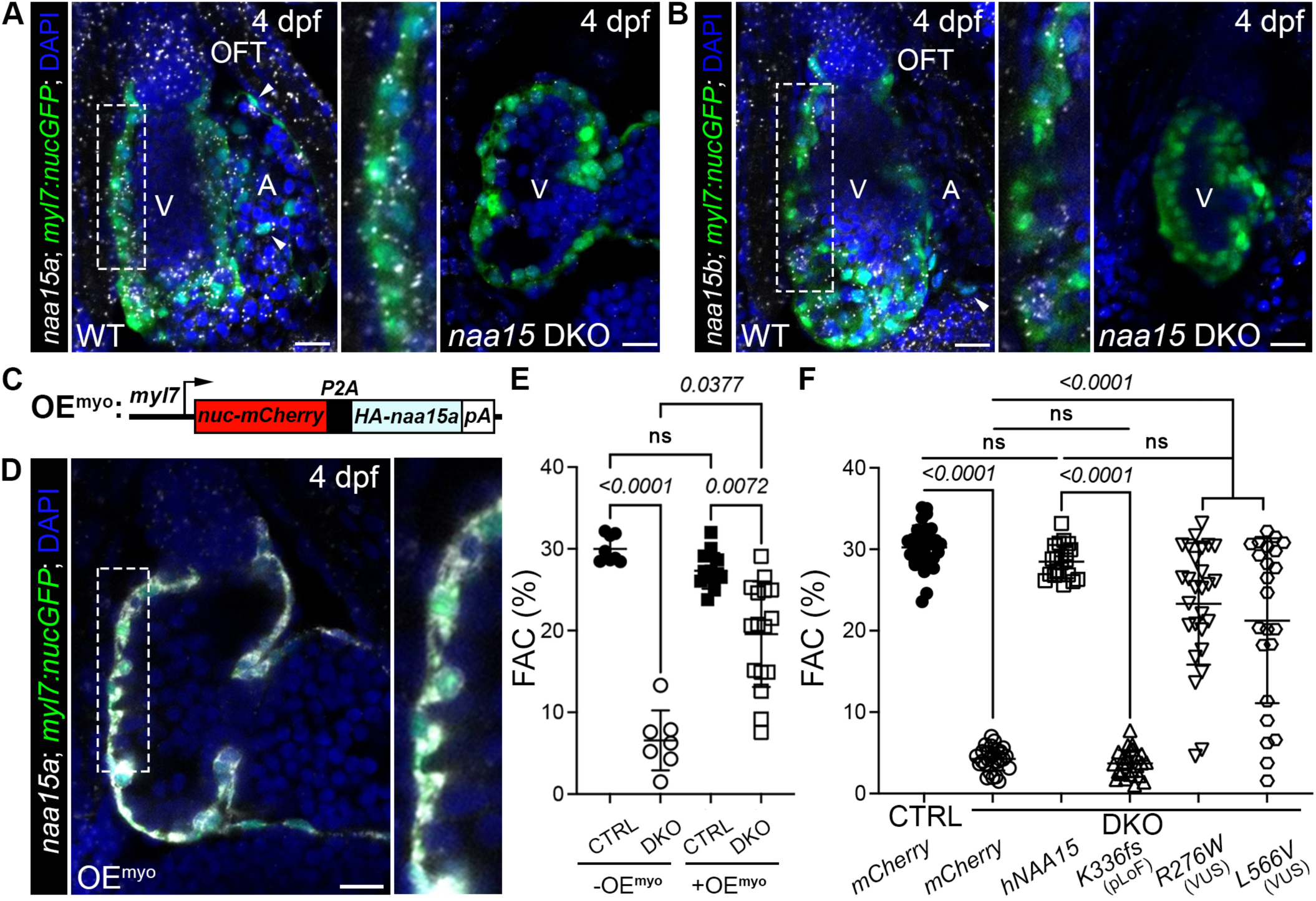
Expression analysis, phenotypic rescue by myocardial *naa15a* re-expression, and human *NAA15* variant testing. (A,B) Single optical sections of hearts in 4 dpf wild-type (WT, left) or double knockout (DKO; right) zebrafish larvae carrying the *myl7:nucGFP* transgene processed for RNAScope in situ hybridization with *naa15a* (A) or *naa15b* (B) probes, immunostained for GFP, and counterstained with DAPI. The boxed regions (left) are shown enlarged (middle). Arrowheads highlight examples of *naa15+* atrial CM nuclei. N=25/25 larvae per group showed the same staining patterns. (C) Schematic diagram of the transgene used to overexpress (OE) Naa15a in the myocardium (OE^myo^). (D) Single optical section of the heart in a 4 dpf OE^myo^ larvae carrying the *myl7:nucGFP* transgene processed for RNAScope in situ hybridization with the *naa15a* probe, immunostained for GFP, and counterstained for DAPI. The boxed region (left) is enlarged (right). N=25/25 larvae per group showed the same staining pattern. (E) Dot plot showing the percent fractional area changes (%FACs) of control-sibling (CTRL) animals without (n=8) or carrying (n=13) the OE^myo^ transgene and DKO animals without (n=7) or carrying (n=16) the OE^myo^ transgene. (F) Dot plot showing the %FACs for 4 dpf CTRL or DKO animals injected at the one-cell stage with mRNAs encoding mCherry, wild-type human NAA15, or the indicated variant-containing human NAA15 isoforms. For (E,F), statistical significance was determined by Kruskal-Wallis Tests followed by a Dunn’s multiple comparisons tests. Adjusted p-values are shown. Means ± 1SD are shown. Scale bars=25 μm. Abbr: V, ventricle; A, atrium; ns, significant; pLoF, predicted loss-of-function; VUS, variant of uncertain significance.

To determine if re-expressing Naa15 exclusively in the myocardium would be sufficient to rescue the DKO ventricular defects (Fig. 1; Fig. 2), we generated a transgenic strain, *Tg(myl7:nuc-mCherry-P2A-HA-naa15a)*, termed “OE^myo^”, designed to overexpress (OE) HA-tagged Naa15a exclusively in the myocardium (myo) from the *myl7* promoter (Fig. 4C)^46^. We confirmed myocardial-specific overexpression of *naa15a* in OE^myo^ animals by RNAScope in situ hybridization at 4 dpf (Fig. 4D). At this stage, OE^myo^ hearts displayed normal %FAC measurements (Fig. 4E) and ventricular sizes (Fig. S6A), demonstrating that myocardial *naa15* overexpression does not affect cardiac contractility or growth. Thereafter, OE^myo^ animals were adult-viable, fertile, and otherwise grossly indistinguishable from WT animals.

DKO animals carrying the OE^myo^ transgene exhibited a partial rescue of %FAC from a baseline of 7% in DKO animals to 20% in DKO + OE^myo^ larvae (Fig. 4E). The rescue fell short of the ∼29% FACs observed in WT larvae, with or without Naa15a overexpression (Fig. 4E). DKO + OE^myo^ hearts also exhibited partial rescue of ventricular size (Fig. S6A). Despite partial rescue across the population, some DKO + OE^myo^ values fell within the control ranges of OE^myo^ animals (6/16 for %FAC and 9/16 for ventricular size; Fig. 4E; Fig S6A), indicating that complete rescue was achieved in a subset of animals. Although the source of variability in rescue is unknown (See discussion), these data demonstrate that *naa15* performs an essential function in the ventricular myocardium by promoting contractility and growth.

### Functional testing of human NAA15 variants in DKO zebrafish

Although the high sequence identity between zebrafish and human NAA15 proteins (Fig. S1; Fig. S2A) suggests they share conserved molecular activities, we tested this hypothesis by injecting one-cell-stage DKO embryos with human *NAA15* mRNA, measuring chamber areas on 4 dpf, and calculating %FACs. Whereas hearts in DKO larvae injected with control mCherry mRNA were unaffected (Fig. 4F; Fig. S6B), DKO animals mis-expressing human *NAA15* showed a complete rescue of both parameters (Fig. 3F; Fig. S6B), demonstrating that NAA15’s molecular function remains exquisitely conserved between species.

Predicted loss-of-function (pLoF) and missense variants of uncertain significance (VUS) in *NAA15* were recently linked to pediatric heart disease^6,10^. To determine whether CHD-linked variants undermine NAA15’s molecular activity, we compared their rescuing activities to that of WT *NAA15*. We injected one-cell-stage DKO embryos with human *NAA15* mRNAs containing variants encoding K336fs (fs, frame shift; pLoF), R267W (VUS), or L566V (VUS), all linked to single-ventricle disease^6^. The truncating K336fs isoform failed to rescue both the DKO contractility and ventricular size phenotypes (Fig. 4F; Fig. 6SB), supporting this variant’s categorization as damaging or LoF^3,6^. The rescuing activities of R276W and L566V mRNAs were variable. Population-wide, they successfully rescued %FAC (Fig. 4F) but not ventricular size (Fig. S6B). However, as with the OE^myo^ rescue, there was significant inter-animal variability. High proportions of hearts exhibited values within the ranges of control animals injected with WT *NAA15* (%FAC: 13/27 for R276W; 11/23 for L566V; ventricular size: 12/27 for R276W; 13/23 for L566V). While the source of the variability is unknown, these data eliminate the possibility that R276W and L566V are LoF variants like K336fs (See discussion).

### Adult naa15^RD^ animals exhibit cardiac defects overlapping those in DKO hearts

Although DKO larvae were inviable beyond 8 dpf, most animals carrying 3 mutant alleles of *naa15* (*naa15a^+/-^*, *naa15b^-/-^*or *naa15a^-/-^*, *naa15b^+/-^* animals) survived to 90 dpf, after which their survival declined steadily, approaching zero by 330 dpf (Fig. 5A). These animals with a reduced genetic <u>d</u>osage of *naa15* (i.e., *naa15^RD^* animals) were confirmed to express significantly less Naa15a and Naa15b protein in their hearts, as determined by deep quantitative proteomic profiling (See below). At 9 months post-fertilization (mpf), *naa15^RD^* animals were diminutive, reflected in reduced body length, weight, and body mass index (BMI; Fig. 5B,C). Like DKO ventricles (Fig. 1B,C; Fig. 2), *naa15^RD^* ventricles were smaller, which held true after normalization to body length (Fig. 5D,E). Histological sectioning and acid fuchsin orange G (AFOG) staining revealed no evidence of widespread scarring (Fig. 5F), suggesting that large-scale apoptosis and replacement fibrosis did not account for the reduced ventricular size. To examine myofibrils, we dissociated 9 mpf WT and *naa15^RD^* hearts to single cells, performed immunostaining for tropomyosin, and binned cells after confocal imaging into categories reflecting “organized” or “disorganized” myofibrils (Fig. 5G). Whereas 100% of CMs from WT hearts contained highly organized myofibrils, that number dropped to 68% in *naa15^RD^* hearts, with the balance of 32% showing the disorganized phenotype (Fig. 5H). Lastly, we measured the size of CMs dissociated from WT and *naa15^RD^* hearts and determined that *naa15^RD^* CMs were smaller (Fig. 5I), consistent with a cellular growth defect. Ultimately, these data suggest that *naa15^RD^* hearts grew less robustly than WT hearts during animal maturation, likely due to molecular and cellular differences similar to but less severe than those in *naa15*-deficient larvae (Fig. 2).

**Figure 5.**
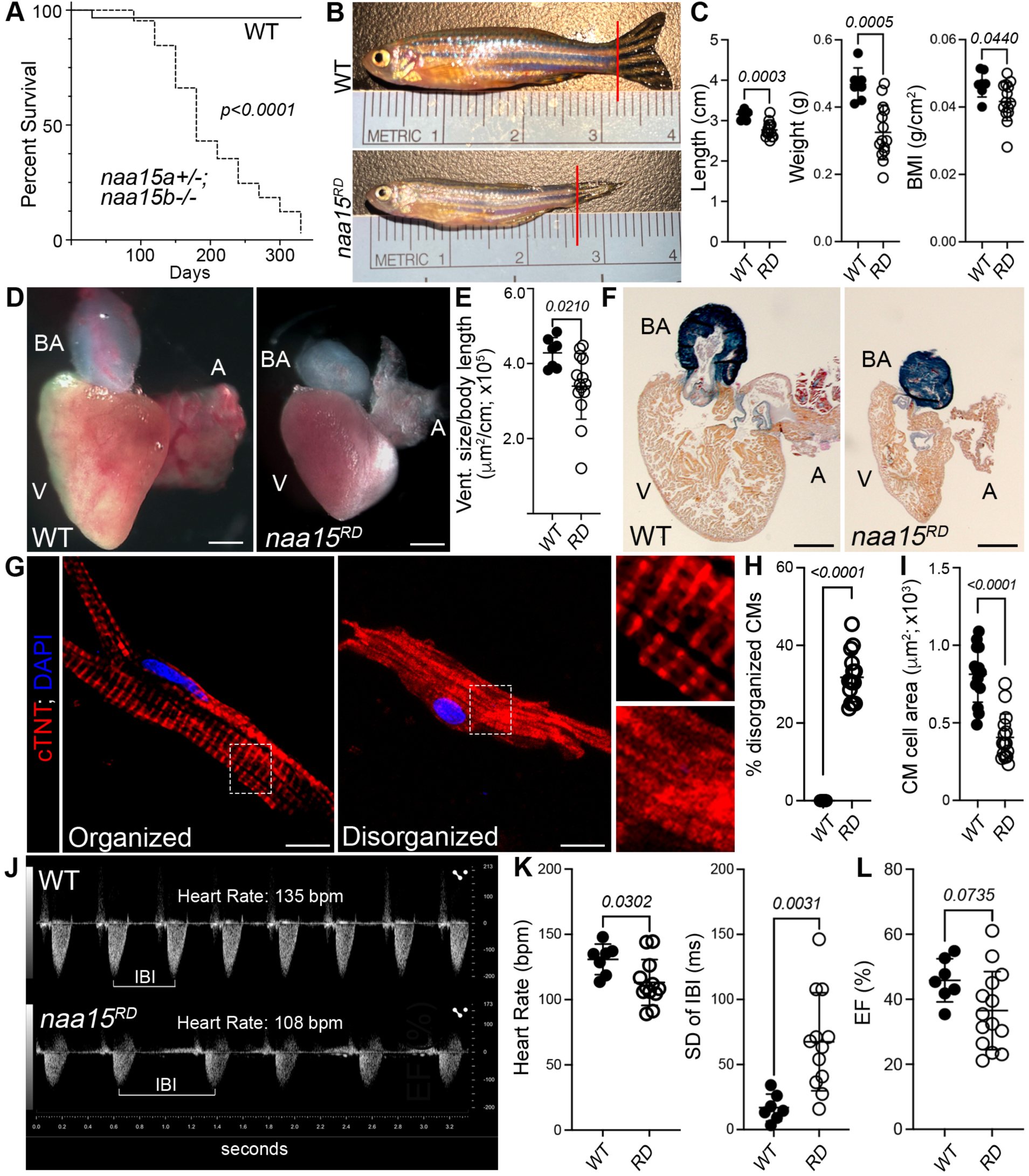
Adult zebrafish with reduced *naa15* dosage exhibit cardiac defects overlapping those in DKO larvae. (A) Kaplan-Meier curve for wild-type (WT; n=60) and *naa15^RD^*(specifically, *naa15a^+/-^*, *naa15b^-/-^*) zebrafish (n=65). Statistical significance was determined by a Log-rank (Mantel-Cox) test. (B) Brightfield images of WT and *naa15^RD^* zebrafish at 9 months post-fertilization (mpf). The red lines show the posterior boundaries of the body length measurements, excluding the fins. (C) Dot plots of body length, weight, and body mass index (BMI) for WT (n=7) and *naa15^RD^* (RD; n=14) zebrafish at 9 mpf. Statistical significance for body length was determined by the Mann-Whitney test. (D) Brightfield images of 9 mpf WT and *naa15^RD^* hearts after dissection. Scale bars=300μm. (E) Dot plot of ventricular areas of 9 mpf WT (n=7) and *naa15^RD^* (n=14) hearts calculated from brightfield images of dissected hearts, examples of which are shown in (D), normalized to body length. (F) Brightfield images of cardiac sections from 9mpf WT (n=4 or more sections from each of 7 hearts) and *naa15^RD^* (n=4 or more sections from each of 14 hearts) animals stained with acid fuchsin orange G (AFOG). No sections exhibited fibrosis. Scale bars=250μm. (G) Confocal images of single cardiomyocytes (CMs) from 9 mpf WT (left) and *naa15^RD^*(middle) hearts immunostained for cardiac Troponin T (cTNT), counterstained with DAPI, and categorized as having organized or disorganized sarcomeres, respectively. Boxed areas are shown at higher magnification (right). Scale bars=10μm. (H) Dot plot showing the percentages of CMs with disarrayed sarcomeres in 9 mpf WT (n=14) and *naa15^RD^* (n=15) hearts. Eighty CMs were examined per heart. (I) Dot plot of dissociated CM cell areas from 9 mpf WT (n=14) and *naa15^RD^* (m=15) hearts. Each data point is the average area of 5-10 CMs from one heart. (J) Representative pulsed-wave Doppler ultrasounds of blood velocity at the bulboventricular junctions of 9 mpf WT and *naa15^RD^* hearts. (K) Dot plots showing heart rates in beats per minute (bpm; left) and standard deviations (SDs) of interbeat intervals (IBIs; right) of 9 mpf WT (n=7) and *naa15^RD^* (n=12) hearts. (K) Dot plot showing the ejection fractions (EFs) of 9 mpf WT (n=7) and *naa15^RD^* (n=14) hearts derived from B-mode ultrasound images shown in Fig. S7. For all dot plots, unless otherwise stated, statistical significance was determined by Student’s t-tests. Means ± 1SD are shown. Abbr: V, Ventricle; A, Atrium; BA, bulbus arteriosus.

Using previously described Doppler echocardiography methods applied to adult zebrafish^50^, we assessed 9 mpf WT and *naa15^RD^*animals for abnormalities in cardiac rhythm and contractility. Hearts in *naa15^RD^*animals beat slower and exhibited greater heart rate variability, as evidenced by an increase in the standard deviation of interbeat intervals (Fig. 5J,K), suggestive of sinus node or autonomic nervous system dysfunction^72,73^. In addition to the reduced heart rates, the stroke volumes of *naa15^RD^* were also lower, resulting in reduced cardiac output (Fig. S7D). However, this reduction in cardiac output disappeared when normalized to ventricular size (Fig. S7E). The ejection fraction, a measure of systolic function^74^, trended lower for *naa15^RD^* hearts but was not statistically significant (Fig. 5L). Ultimately, these data demonstrate that adult *naa15^RD^* hearts showed phenotypic similarities to DKO hearts, including diminutive ventricles with reduced heart rates, smaller CMs with disorganized sarcomeres, and a functional parameter that trended lower but did not reach statistical significance.

### Deep quantitative proteomic profiling uncovers mitochondrial protein dysregulation in naa15^RD^ adult hearts

NatA is responsible for N-terminal acetylation of large subsets of proteins in diverse organisms from yeast to humans^16^. Because one function of this modification is to regulate protein stability, we sought to identify candidate NatA targets among proteins differentially expressed in DKO hearts, as revealed by proteomics analysis. Initially, we planned to perform deep quantitative proteomic profiling of WT and DKO hearts before 8 dpf. However, we determined that the number of hearts per replicate for proteomics analysis was prohibitively high based on our empirical protein yield. Because of the phenotypic overlap between *naa15*-deficient larval hearts and *naa15^RD^* adult hearts, as an alternative, we performed proteomic profiling of adult hearts from WT and *naa15^RD^*animals, reasoning that both genotypes likely share qualitatively similar proteomic disturbances. Moreover, given that CHD probands were heterozygous for *NAA15* variants^6^, the proteomic alterations in *naa15^RD^* hearts might be a better reflection of the human condition.

We extracted total protein from dissected ventricles of 9 mpf WT and *naa15^RD^* animals (genotype *naa15a^+/-^, naa15b^-/I^*^-^) after pooling 3-5 ventricles in each of 5 replicates per group. We performed LysC and Trypsin digestion, tandem mass tag (TMT) 16 isobaric labeling, reverse phase proteome fractionation, and liquid chromatography tandem mass-spectrometry (LC-MS/MS; Fig. 6A). We detected 9514 total proteins, 9431 of which were considered fully quantified based on their presence in 70% of replicates (Table 1). Principal component analysis showed replicate segregation by experimental group (Table 1). We identified differential expression of 3593 proteins in *naa15^RD^* hearts (p-adj>0.05), with 1910 and 1683 showing decreased and increased expression, respectively (Fig. 6B; Table 1). As anticipated, both Naa15a (log_2_FC= –5.8) and Naa15b (log_2_FC= –15.3) were significantly decreased, validating *naa15^RD^* animals as a model of reduced Naa15 dosage. Other members of NatA and NatE, such as Naa10 (Log_2_FC= –8.7) and Naa50 (Log_2_FC= –5,4), were also less abundant, as was the NatA-interacting protein Hypk (Log_2_FC= –4.3), demonstrating that several members of Naa15-containing complexes were destabilized in adult *naa15^RD^*hearts. The observation that Naa10 and Hypk protein levels were reduced in *naa15^RD^*adults, while their transcript levels were increased in DKO larvae (Fig. S2), is consistent with cellular attempts at compensation in DKO larvae.

**Figure 6.**
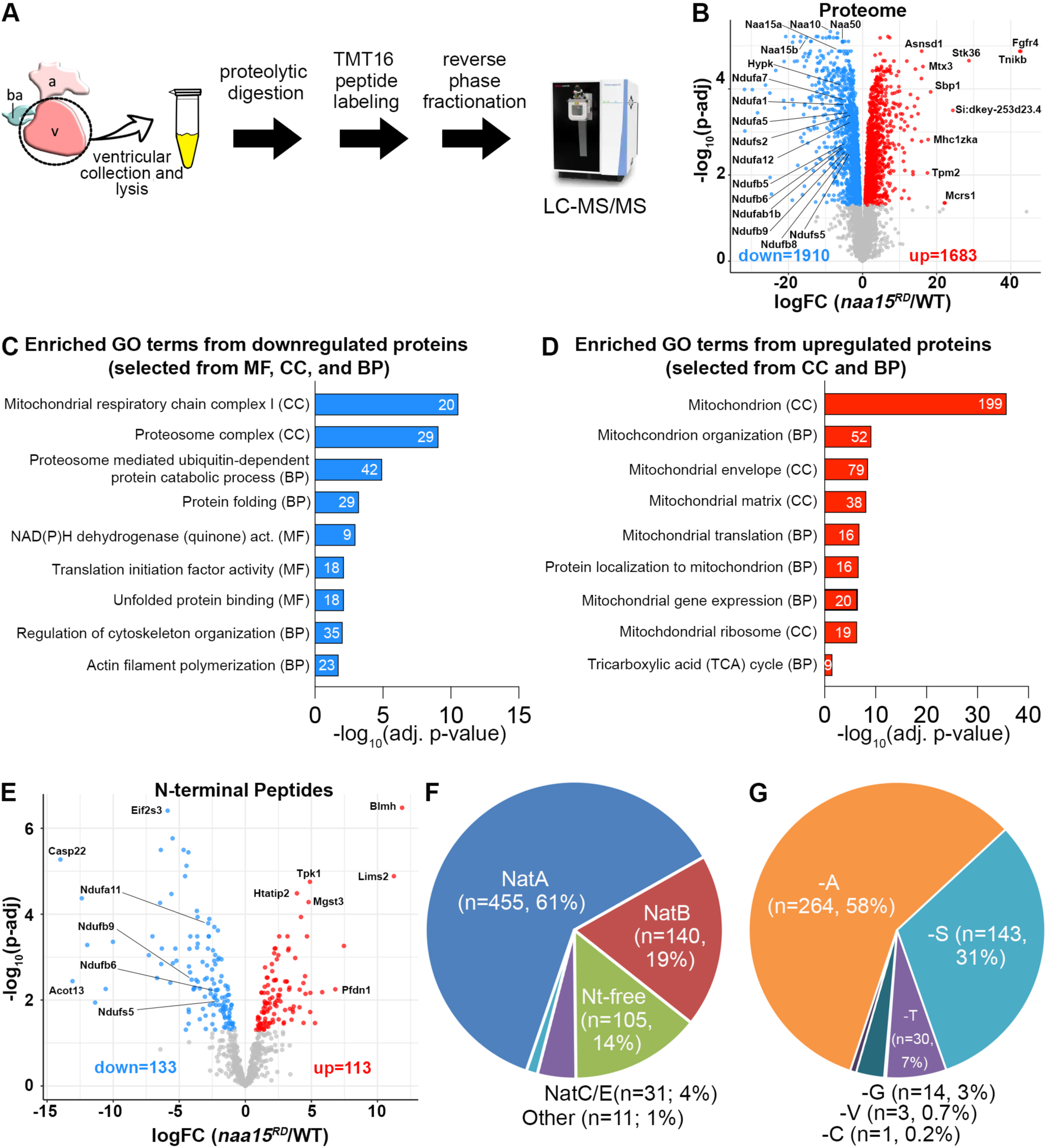
Deep quantitative proteomic profiling of adult *naa15^RD^* hearts. (A) Schematic diagram of the deep quantitative proteomics profiling workflow performed on 9 months post-fertilization (mpf) wild-type (WT) and *naa15^RD^* ventricles. (B) Volcano plot showing fold changes (FCs; expressed as log_2_FCs) and adjusted p-values [p-adj; expressed as -log10(p-adj)] of differentially expressed peptides between 9 mpf WT (n=5 biological replicates) and *naa15^RD^* ventricles (n=5 biological replicates). Peptides meeting the inclusion criteria [adjusted p-value <0.05] for the gene ontology (GO) term enrichment analysis are shown in blue or red. NatA subunits and interacting proteins, as well as several mitochondrial respiratory complex I subunits are highlighted in the downregulated proteins. (C,D) Bar graphs showing selected GO terms enriched from three GO categories Molecular Function (MF), Cellular Component (CC), and Biological Process (BP) in the down (C) and upregulated (D) proteins in *naa15^RD^* ventricles. (E) Volcano plot showing fold changes (expressed as log_2_FCs) and adjusted p-values [expressed as -log10(p-adj)] of differentially expressed N-terminal peptides between 9 mpf WT (n=5 biological replicates) and *naa15^RD^* ventricles (n=5 biological replicates). Peptides meeting inclusion criteria (adjusted p-value <0.05) for the GO term enrichment analysis are shown in blue or red. N-terminal peptides of four subunits of the mitochondrial respiratory complex I are highlighted among the downregulated proteins. (F) Pie chart showing the absolute numbers and percentages of fully quantified N-terminal peptides detected in our dataset that were unacetylated (Nt-free) or acetylated by the indicated Nat complex. (G) Pie chart showing the absolute numbers and percentages of fully quantified N-terminal peptides with the indicated amino acids in the second position acetylated by NatA after initiator methionine removal.

To determine if the differentially expressed proteins share common features or functions, we performed Gene Ontology (GO) term enrichment analysis. Among the downregulated proteins, we identified several similar or related GO terms enriched across two or more GO categories (Table 2). For example, the terms “Mitochondrial respiratory chain Complex I” and “NAD(P)H dehydrogenase (quinone) activity” were the top terms in the Cellular Component (CC) and Molecular Function (MF) categories based on adjusted p-values (Fig. 6C; Table 2). This reflected reduced expression of 20 subunits of respiratory complex I of the mitochondrial electron transport chain (ETC)^75^. During oxidative phosphorylation, four ETC protein complexes (I-IV) perform sequential redox reactions to establish a proton gradient across the inner mitochondrial membrane, which drives ATP production by ATP synthase as protons move back down the gradient. Respiratory complex I transfers electrons from NADH to ubiquinone while pumping electrons into the intermembrane space from the mitochondrial matrix. Complex I activity also generates reactive oxygen species (ROS), which can modulate several downstream signaling pathways^75,76^. Disruptions in complex I activity compromise ATP production, increase ROS production, and are associated with several human diseases^75,77^.

**Table 2.** Spreadsheets showing the Gene Ontology terms enriched in the gene sets encoding differentially expressed proteins (DEPs) that were up- or down-regulated in *naa15^RD^* hearts (adjusted p<0.05, FC>|1|) within the Biological Process (BP), Cellular Component (CC), or Molecular Function (MF) categories.

The proteomics analysis also highlighted reduced expression of factors required for proteome homeostasis, including those involved in translation initiation, protein folding, unfolded protein binding, and ubiquitin-dependent proteosomal degradation (Fig. 6C; Table 2). Lastly, it uncovered lowered expression of proteins involved in cytoskeletal organization and actin filament polymerization, consistent with the documented sarcomere defects in a subset of *naa15^RD^* CMs (Fig. 5G, H; Fig. 6C; Table 2).

Among the upregulated proteins, we observed a predominance of GO terms related to mitochondria, including those pertaining to mitochondrial structure, translation, gene expression, and metabolism (Fig. 6D; Table 2). All in all, the GO term analysis suggests that reduced Naa15 activity compromises the expression of several protein classes, including numerous subunits of the ETC’s respiratory complex I, which would undermine mitochondrial function. It also stimulates upregulation of multiple proteins localized to or functioning within mitochondria, consistent with a compensatory response.

### Cross-species comparisons of differentially expressed proteins downstream of NAA15

Next, we determined whether significant overlap exists between zebrafish and human protein orthologs that were differentially expressed in *NAA15^RD^*adult zebrafish hearts and *NAA15^+/-^* or *NAA15^-/-^* undifferentiated human iPSCs^6^. To that end, we identified human orthologs of the proteins fully quantified in *naa15^RD^* hearts (Table 1; Materials and Methods), cataloged protein pairs dysregulated in both species, and performed Fisher’s exact tests to assess the statistical significance of shared proteomic disturbances. Pairwise comparisons between the zebrafish dataset and the human stem cell datasets from heterozygous or null cells revealed some overlap but not enough to reach statistical significance (Table 5; Figure S8). Accordingly, our GO term analysis did not uncover enrichment of terms related to cellular ribosomes, which was a predominant signal in undifferentiated human iPSCs genetically engineered to contain reduced or absent NAA15 activity^6^. Targeted searches for the zebrafish orthologs of the four CHD proteins reduced in *NAA15* heterozygous or null iPSCs^6^ revealed no detection (DHCR7 and NSD1) or no differential expression (MAP2K2 and RPL5) in *naa15^RD^* hearts (Table 5). Ultimately, the cross-species comparison suggests that the direct and indirect targets of *NAA15,* as a whole, are context-dependent (See discussion).

### Characterization of the N-terminomes of WT and naa15^RD^ adult zebrafish hearts

Next, we identified N-terminal peptides within the proteomic datasets of WT and *naa15^RD^* adult hearts. We detected 1057 previously annotated N-terminal peptides, 742 of which were fully quantified across 70% of the replicates (Fig. 6E; Table 3). Similar numbers of N-terminal peptides were detected in undifferentiated human iPSCs^6^. Most of the peptides were N-terminally acetylated (637 of 742 or 86%; Fig. 6F; Table 3). Among these, 455 (61%) were NatA targets, based on the absence of an initiation methionine and Nt-acetylation of amino acids A, S, T, C, G, or V (Fig. 6F)^16^. Acetylation occurred most frequently on Alanine (58%) or Serine (32%), consistent with a previous analysis (Fig. 6G)^6^. Smaller percentages were acetylated on Threonine (7%), Glycine (3%), Valine (0.7%), or Cysteine (0.2%; Fig. 6G). Additional acetylated N-terminal peptides were identified as NatB (140, 19%), NatC (30, 4%), or NatE (1, 0.1%) targets based on known substrate specificities (Fig. 6F)^16^. Lastly, 105 N-terminal peptides (14%) lacked N-terminal acetylation (Fig. 6F) even though this group contained potential NatA (24), NatB (10), NatC (30), and NatA/E (3) targets based on protein sequence. Only two N-terminal peptides, including one NatA target, were detected in both acetylated and non-acetylated forms, suggesting that non-acetylated Nat targets become susceptible to degradation, consistent with previous observations^16^.

**Table 3.** Spreadsheets showing the fully quantified N-terminal peptides and differentially expressed N-terminal peptides (DENtPs; adjusted p<0.05, FC>|1|) in *naa15^RD^*hearts as determined by deep quantitative proteomic profiling. NatA substrates are highlighted in separate sheets. The log_2_FC values are reported as both WT/*naa15^RD^*and *naa15^RD^*/WT (gray background).

Between WT and *naa15^RD^* hearts, 246 N-terminal peptides showed differential expression (p-adj<0.05), 133 of which were down-regulated and 113 were up-regulated in *naa15^RD^* hearts (Fig. 6E). No enriched GO terms were identified among the down-regulated N-terminal peptides based on adjusted p<0.05. Using an unadjusted p<0.05, ‘mitochondrial protein containing complex’ (CC) was enriched, based, in part, on the downregulation of N-terminal peptides from the ETC’s respiratory complex I subunits Ndufb6, Ndufb9, Ndufa11, and Ndufs5 (Fig. 6E), the first three of which are NatA targets detected in acetylated forms (Table 4). Among the up-regulated N-terminal peptides, ‘mitochondrion’ (CC) was enriched based on adjusted p<0.05. Using an unadjusted p<0.05, additional terms were enriched, including “mitochondrial organization” (BP) and ‘mitochondrial membrane’ (CC). Among the 246 differentially expressed acetylated N-terminal peptides, 142 (57%) were NatA targets, 94 of which showed downregulation, consistent with reduced NatA activity. Another 48 exhibited upregulation, suggesting that compensatory mechanisms can stabilize some NatA targets or that N-terminal acetylation directly or indirectly destabilizes a subset of targets in WT hearts^16^.

**Table 4.** Spreadsheets showing the Gene Ontology terms enriched in the gene sets encoding proteins with differentially expressed N-terminal peptides (DENtPs) that were up- or down-regulated in *naa15^RD^*hearts (adjusted p<0.05, FC>|1|) within the Biological Process (BP), Cellular Component (CC), or Molecular Function (MF) categories.

Lastly, we manually cross-referenced the 34 human proteins that showed differential N-terminal acetylation in *NAA15* haploinsufficient or null iPSCs^6^ with the differentially expressed N-terminal peptides in *naa15^RD^* zebrafish hearts. Among the 34 human proteins, we identified high-confidence zebrafish orthologs for 30. We detected N-terminal peptides for 6 in adult zebrafish hearts, with one showing a significant reduction in *naa15^RD^* hearts, the N-terminally acetylated NatA target Ndufb6 of respiratory complex I (Table 3; Table 5). Ultimately, the N-terminal acetylation defects documented in zebrafish hearts and undifferentiated human iPSCs with compromised *NAA15* activity were almost entirely distinct, which we attribute to context-dependent NatA functions (See discussion).

**Table 5.** Spreadsheets showing differential expression values for 2013 orthology pairs in zebrafish *naa15RD* hearts or *NAA15* haploinsufficient or null human induced pluripotent stem cells relative to control samples, including the results of Fisher’s exact tests. Also included is a cross-species comparison of dysregulated N-terminal acetylation events between the same experimental groups.

### Mitochondrial morphology and functional defects in DKO larvae

Given the prominent mitochondrial-related signals in the GO term analysis of *naa15^RD^* hearts, we hypothesized that *naa15*-deficient myocardium would exhibit abnormal mitochondrial density, size, or content as evaluated by ultrastructural imaging. Therefore, we performed transmission electron microscopy on 4 dpf WT and DKO animals. Evaluating multiple sections from three hearts each, we identified and outlined CMs based on the presence of intracellular muscle fibers and cell boundaries. We measured the total CM area per section, the density of mitochondria within that area, the area of each mitochondrion, and the total mitochondrial area divided by total CM area. Expressing these measurements as ratios, we found that DKO CMs had less dense, smaller mitochondria, which equated to reduced total CM mitochondrial content (Fig. 7A-E).

**Figure 7.**
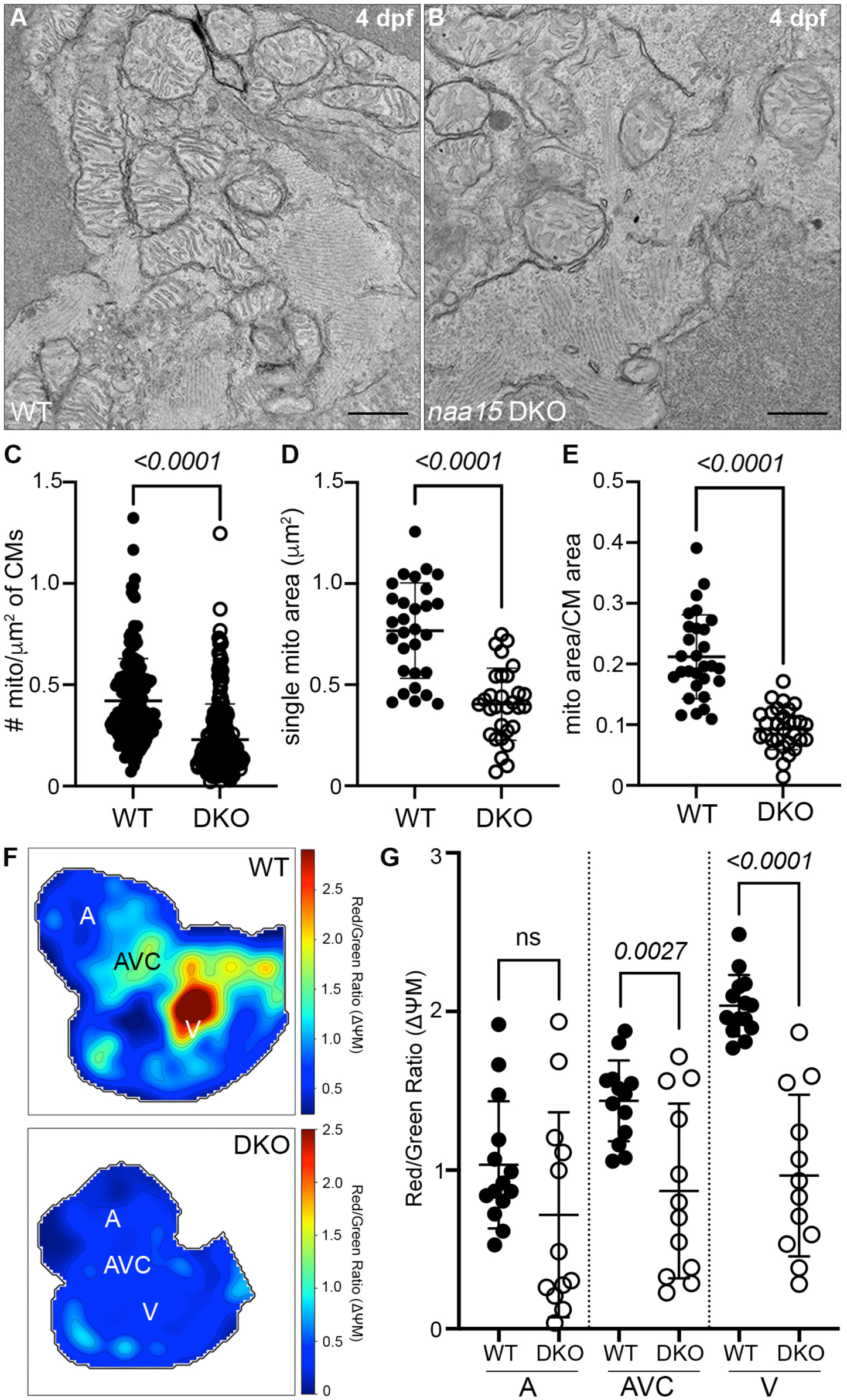
Reduced mitochondrial content, size, and membrane potential in *naa15*-deficient hearts. (A,B) Transmission electron micrographs of wild-type (WT) and double knockout (DKO) hearts on 4 days post-fertilization (dpf). Scale bars=500nm. (C-E) Dot plots showing the numbers of mitochondria per cardiomyocyte (CM) area (μm^2^; C), individual mitochondrial areas (μm^2^; D), and ratios between the total mitochondrial and CM areas (E) measured in transmission electron micrographs from 4 dpf WT and DKO hearts. For WT hearts, n=29 total micrographs (9, 10, and 10 across 3 hearts). For DKO hearts, n=30 total micrographs (10,10, and 10 across 3 hearts). For individual mitochondrial areas, n=6-8 mitochondria were measured from the same micrographs. n=188 total mitochondria for WT and n=222 mitochondria for DKO hearts. (F) Heat maps showing the ratios of red to green fluorescence intensities in representative 4 dpf WT and KO hearts. (G) Dot plot showing the ratios of red to green fluorescence in the atria (A), atrioventricular canals (AVC), and ventricles of 4 dpf WT (n=14 for A and V, n=13 for AVC) and DKO (n=12) hearts. Statistical significance was determined by Student’s t-tests (C,D,F) or a Mann-Whitney test (E). Means ± 1SD are shown.

Because disruptions in mitochondrial respiratory complex I activity would be predicted to depolarize the mitochondrial membrane, we evaluated relative membrane potentials between WT and *naa15*-deficient larval hearts with optical mapping after exposure to JC-1 dye^78^. JC-1 exists in a monomeric form that emits green fluorescence when distributed at lower concentrations throughout the cell. In WT cells, the accumulation of JC-1 in the negatively charged mitochondrial matrix induces the formation of J-aggregates, which fluoresce red. In unhealthy mitochondria with decreased membrane potential and increased matrix charge, JC-1 accumulation is attenuated, resulting in fewer aggregates and a lower red signal. Therefore, the ratio of red to green fluorescence intensities can serve as a relative indicator of mitochondrial membrane potential between two experimental conditions. When we compared the red to green ratios between WT and *naa15*-deficient larval hearts, we observed a significant ∼50% reduction in *naa15*-deficient ventricles (Fig. 7F,G). This reduction was also observed at the atrioventricular canal but not the atrium, despite a trend towards reduced potential. These data demonstrate that *naa15* activity is indispensable for establishing or maintaining mitochondrial membrane potential and function in ventricular myocardium.

## Discussion

We report the first vertebrate animal model with deficient or reduced *NAA15* activity that exhibits cardiac abnormalities. Zebrafish larvae lacking *naa15a* and *naa15b*, both expressed in the myocardium, showed reductions in ventricular size, contractility, and heart rate before death at approximately one week post-fertilization. The smaller ventricles were attributable to fewer diminutive CMs that were incapable of proliferation. On a subcellular level, DKO CMs exhibit mild myofibril disarray and reduced mitochondrial content and function. Ventricular contractility and growth defects were partially rescued by CM-specific overexpression of zebrafish *naa15a* and fully rescued by ubiquitous mis-expression of human *NAA15*, thereby enabling functional testing of human *NAA15* variants. Outside of the heart, the lower jaws of DKO animals were underdeveloped and associated with abnormal growth or positioning of pharyngeal cartilage and muscles. Adult zebrafish carrying a single functional copy of *naa15*, of four total, survived up to one year. These animals with a reduced genetic dose of *naa15* (*naa15^RD^*) exhibited cardiac phenotypes that overlapped with those of DKO larvae, including smaller ventricles, reduced heart rates, and diminutive CMs with evidence of myofibril disorganization. A trend towards reduced systolic function fell short of statistical significance.

A previous study evaluated the expression patterns and knockdown phenotypes of *naa15a* and *naa15b* in zebrafish^19^, but no information was provided for the heart. Morpholino knockdown of *naa15a* and *naa15b* caused most embryos to die before 24 hpf. The surviving embryos were diminutive with body curvature and myotome defects. Cardiac abnormalities were not reported. By contrast, DKO larvae were normally sized without body curvature defects, and they exhibited multiple cardiac defects by 4 dpf. Discrepancies between morphant and mutant phenotypes are common and attributable to morpholino off-target effects, generalized morpholino toxicity, or transcriptional adaptation in mutants^43,79–82^. Alternatively, morphant-specific phenotypes could be attributable to translational inhibition of maternally deposited *naa15a* and *naa15b*. In future experiments, it could be of interest to determine whether *naa15* mutants exhibit myotome defects as reported in morphants^19^, or evidence of neurodevelopmental phenotypes, as described in human *NAA15*-related syndrome^9^.

We anticipate that *NAA15* performs a conserved and indispensable role in cardiac development across the animal kingdom. This likelihood is supported by our zebrafish data, the phenotypes of haploinsufficient and null *NAA15* iPSC-CMs^6^, and those of *Drosophila* with cardiac-specific silencing of the *NAA15* ortholog^25^. Additional support comes from developmental expression studies of *NAA15* orthologs, revealing conserved localization to embryonic hearts in flies, birds, and several mammalian species^25,83,84^. A mouse model of *NAA15*-related heart disease would be a valuable resource for bridging observations between zebrafish and human cell culture. To that end, *Naa15* null mice were recently reported to be embryonic lethal, consistent with a potential CHD phenotype, but the cause and timing of death were not reported^15^.

The phenotypes of *NAA15*-null human iPSCs^6^ and *Drosophila* with cardiac-specific *NAA15* silencing^25^ suggest that *NAA15* is required for CM differentiation during heart development. Specifically, *NAA15*-null human iPSCs failed to assume CM fates by directed differentiation^6^ and a fly strain with cardiac-specific knockdown of the *NAA15* ortholog was incapable of forming a heart^25^. By contrast, DKO zebrafish were clearly capable of CM differentiation. These disparate observations could reflect species-specific requirements for *NAA15*. More likely, however, is that maternal deposition of *naa15a* and *naa15b* in DKO zygotic mutants enables CM differentiation before their depletion. Determining whether maternal *naa15a* and *naa15b* are required for CM differentiation in zebrafish would require generating maternal-zygotic mutant embryos, which is onerous when null females die before sexual maturity, as is the case for *naa15* DKO females.

*NAA15*’s function in the heart likely extends beyond development, as evidenced by the cardiac phenotypes observed in *naa15^RD^* adults and documented cardiac expression in adults of several higher vertebrate species, including humans^84,85^. Although adult humans express *NAA15* relatively ubiquitously, heart muscle is among the cell types that show prominent expression in the Single Cell Type Atlas^86^ (www.proteinatlas.org; v24). Conditional knockout of *Naa15* in the adult zebrafish or mouse myocardium would unequivocally determine its intrinsic requirement for cardiac homeostasis or function in adulthood.

The ventricular contractility and growth phenotypes in DKO animals were partially rescued by myocardial re-expression of *naa15a*. Nonetheless, approximately one-third (%FAC) to one-half (ventricular size) of the hearts were fully rescued, with their values falling within control ranges. There could be multiple reasons for the heterogeneous rescue. It could arise from suboptimal protein activity due to the HA tag or reduced expression in some animals due to *naa15a’s* downstream position in the bicistronic construct^87^. Alternatively, Naa15 could function in non-myocardial cell types to support ventricular size and contractility, since ubiquitous mis-expression of human *NAA15*, conferred by RNA injection, provided superior rescue compared with myocardial-specific re-expression. Lastly, unknown genetic or epigenetic modifiers could have influenced rescue efficiency. Supraphysiologic *naa15a* expression levels were unlikely to account for the incomplete rescue, as WT ventricles overexpressing *naa15a* were unaffected. Although the source of variable rescue is unclear, our data demonstrate that *naa15* functions in the myocardium to support ventricular contractility and growth.

Ubiquitous mis-expression of human *NAA15* from injected mRNA was sufficient to achieve a complete rescue of DKO hearts. This observation highlights the exquisite conservation of NAA15’s molecular activity, which enabled functional testing of human variants. Mis-expressing the truncated K336fs protein in DKO animals failed to rescue ventricular contractility and size, confirming the absence of molecular activity and the underlying variant’s designation as damaging or LoF^3,6^. Mis-expressing R276W or L556V rescued contractility but not ventricular size across the injected population. While this indicates that the cardiac functional and growth phenotypes are, to some degree, separable, we found positive correlations between these variables in both cohorts (Spearman r=0.71, 95% CI=0.45-0.86, p<0.0001 for R276W-injected animals; Spearman r=0.61, 95% CI=0.25-0.82, p=0.002 for L556V-injected animals), demonstrating that larger hearts have higher %FACS.

For both missense variants, there was significant animal-to-animal variability in the degree of rescue. Approximately half of the data points fell within the control ranges, precluding LoF designations, while the remainder indicated intermediate or no rescue. An incomplete rescue was unlikely due to an inconsistent injection technique, as uniform rescue was observed when the same experimentalist injected WT *NAA15*. Nonetheless, the amount of mRNA injected per embryo by one individual can vary, so it’s plausible that the degree of rescue was sensitive to minor variations in dose, unlike for human *NAA15*. The observation that approximately half of the animals showed complete rescue suggests that the missense variants are “benign”. For R276W, this is consistent with its ability to rescue the growth defect in a yeast model of NatA deficiency^6^. However, given the non-uniform nature of rescue and their rarity in the general population^6^, they could also be hypomorphic. Regardless, this study, as well as others^36^, demonstrates the facile and expeditious nature of variant testing in zebrafish.

Several phenotypes in *naa15*-deficient zebrafish hearts have been associated with or implicated in CHD pathogenesis, including altered hemodynamics^88^, compromised CM cell cycle progression^89^, and mitochondrial dysfunction^90^. Specifically, in single-ventricle disease, mitochondrial phenotypes were documented in the *Ohia* mouse model of HLHS, in primary tissue from HLHS fetuses, and in HLHS patient-derived iPSC-CMs. They included reduced mitochondrial gene expression, membrane potential, size, and content^91–93^. Additional shared phenotypes between *naa15*-deficient hearts and HLHS models include decreased CM contractility, impaired cell cycle progression, and disorganized or reduced myofibrils^92–95^. Most of these phenotypes were also observed in genetically engineered zebrafish or fly models of HLHS, both of which were based on sequencing studies of affected families^36,96^.

There is mounting evidence that mitochondrial defects can be a primary driver of HLHS phenotypes or subsequent heart failure after surgical correction. Specifically, a *Drosophila* model of HLHS relies on knocking down a subunit of the mitochondrial MICOS complex, which was found mutated in a proband with HLHS and is required for normal cristae structure and ATP production^96^. Moreover, disrupting MICOS activity in human iPSC-CMs was sufficient to cause HLHS-related phenotypes, including reduced oxygen consumption, CM proliferation, and filamentous actin. Lastly, sildenafil-mediated restoration of the mitochondrial membrane potential in patient-derived iPSC-CMs rescued several HLHS or heart failure-related cellular phenotypes^93^. These observations have raised the possibility of preventing HLHS-associated heart failure with mitochondrial transplantation^97^.

The proteomic alterations in *naa15^RD^* adult hearts did not significantly overlap with those of haploinsufficient or null *NAA15* human iPSCs^6^. This is perhaps not surprising since the profiled material comprised different cell types (reprogrammed stem cells vs. whole hearts) from different species (human vs. zebrafish) and life stages (embryonic-like vs. adult). Moreover, reduced expression of Naa15 in adult zebrafish resulted in the differential expression of many more NatA-targeted N-terminal peptides (142) than were altered in *NAA15* haploinsufficient or null human stem cells (34)^6^. This difference could result from the presence of an *NAA15* paralogue, *NAA16*, in humans, which is absent in zebrafish^98^.

In the *naa15^RD^* proteomics dataset, we found downregulation of 20 subunits (of 43 total) making up the ‘mitochondrial respiratory chain complex I’. Among these, we identified 2 NatA targets, Ndufb6 and Ndufb9, whose expression levels decreased, including their acetylated N-terminal peptides. A third NatA target, Ndufa11, trended towards reduced expression levels (p=0.077), while its acetylated N-terminal peptide was significantly reduced. Orthologous proteins were found to be N-terminally acetylated in bovine hearts^99^, and they are indispensable for respiratory complex I assembly and function in human cells^100,101^. All three proteins localize to the hydrophobic membrane arm of complex I embedded in the inner mitochondrial membrane^102^. Since N-terminal acetylation increases hydrophobicity, influencing protein stability, folding, subcellular localization, and physical interactions^16^, we speculate that compromised N-terminal acetylation alters these proteins’ incorporation or functioning within the complex. Removing one subunit from the complex often destabilizes other subunits^100^, which could explain why non-NatA targets within complex I also exhibit reduced expression in *naa15^RD^*hearts.

Reductions in membrane potential caused by disruptions in complex I are known to lower ATP production^75^, which would have adverse consequences for ATP-dependent cellular processes, including CM contractility^103^ and proliferation^104^, both of which were documented in DKO hearts. Disruptions in complex I are also known to increase ROS, which can have pleiotropic effects on signaling pathways and cellular physiology^75^. While it is tempting to speculate that respiratory complex I dysfunction is solely or primarily responsible for the documented heart phenotypes in zebrafish, a more conservative interpretation is that disruptions in complex I are among several molecular contributors, given the large number of proteins we found to be modified by NatA and differentially expressed in *naa15^RD^*hearts. More likely, the cardiac phenotypes arise from the combinatorial consequences of reduced N-terminal acetylation across the proteome, inclusive of complex I dysfunction, which together reduce cardiac contractility and cause a hypoplastic ventricle. As such, additional studies on this and other animal or cell-based models will further clarify the molecular mechanisms underlying *NAA15*-related heart disease.

## Acknowledgements

Sanger sequencing, next-generation amplicon sequencing, and microsatellite analysis were performed by the Massachusetts General Hospital DNA Core. Several monoclonal antibodies were obtained from the Developmental Studies Hybridoma Bank, created by the NICHD of the NIH and maintained at the University of Iowa, Department of Biology, Iowa City, IA 52242. DNA oligonucleotides were obtained from Integrated DNA Technologies. Transmission electron microscopy was performed at the Harvard Medical School Electron Microscopy Core Facility. Hui-Min Yin, PhD, generously provided the *pTol2 myl7:nuc-mCherryP2A* backbone for generating the OE^myo^ transgene. We thank Vassilios Bezzerides, MD, PhD, for valuable discussions on NatA functions in the heart.

## Sources of Funding

WPP was supported by a post-doctoral fellowship (22POST903575) from the American Heart Association (AHA). OW was supported by a National Institutes of Health (NIH) training grant (T32HL007572) to Boston Children’s Hospital (PI: WT Pu) and a post-doctoral fellowship (24POST1192144) from the AHA. Alexander Akerberg was supported by a post-doctoral fellowship (20POST35110027) from the AHA. RZ and KC were supported, in part, by K99HG013662 (RZ), R01GM138407 (KC), and R01HL174928 (KC). Research in the Burns Laboratory was supported by NIH grants R35HL135831 (PI:CEB), R01HL171206 (PI:CEB), R01HL176663 (PI:CGB), U01AA030185 (MPI:CEB and CGB), and funds from Boston Children’s Hospital Department of Cardiology.

## Disclosures

The authors have nothing to disclose.

## Author Contributions

WPP designed and performed the majority of experiments, interpreted data, and acquired funding; OW aided in echocardiography data acquisition and analysis; AA and MM isolated the *naa15a^chb13^* and *naa15b^chb14^* alleles and performed early phenotyping; JG and AB aided data acquisition; HRM performed the optical mapping with oversight from CAM; RZ performed the GO term and cross-species analyses with oversight by KC; HK and PH performed the deep proteomic profiling of *naa15^RD^* hearts with oversight by SC; CEB and CGB conceived and supervised the study, designed experiments, interpreted data, and acquired funding; WPP, CEB, and CGB wrote the manuscript with input from all authors.

## Data Availability

The original mass spectra and the protein sequence database used for searches have been deposited in the public proteomics repository MassIVE (http://massive.ucsd.edu) and are accessible at ftp://MSV000101582@massive-ftp.ucsd.edu when providing the dataset password: BurnsNAA15. If requested, also provide the username: MSV000101582. These datasets will be made public upon acceptance of the manuscript.

## Supplemental Figure Legends

**Figure S1.**
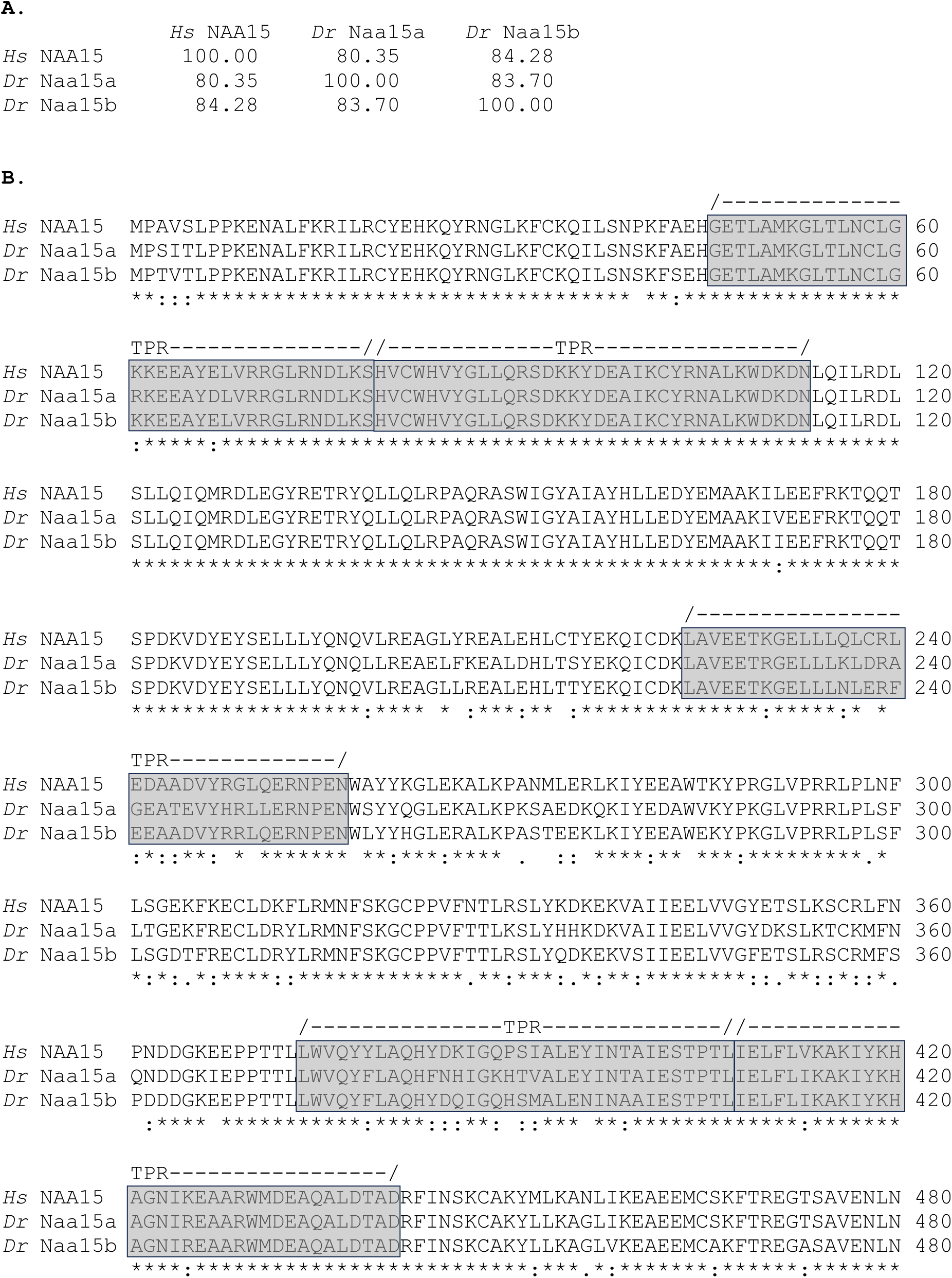
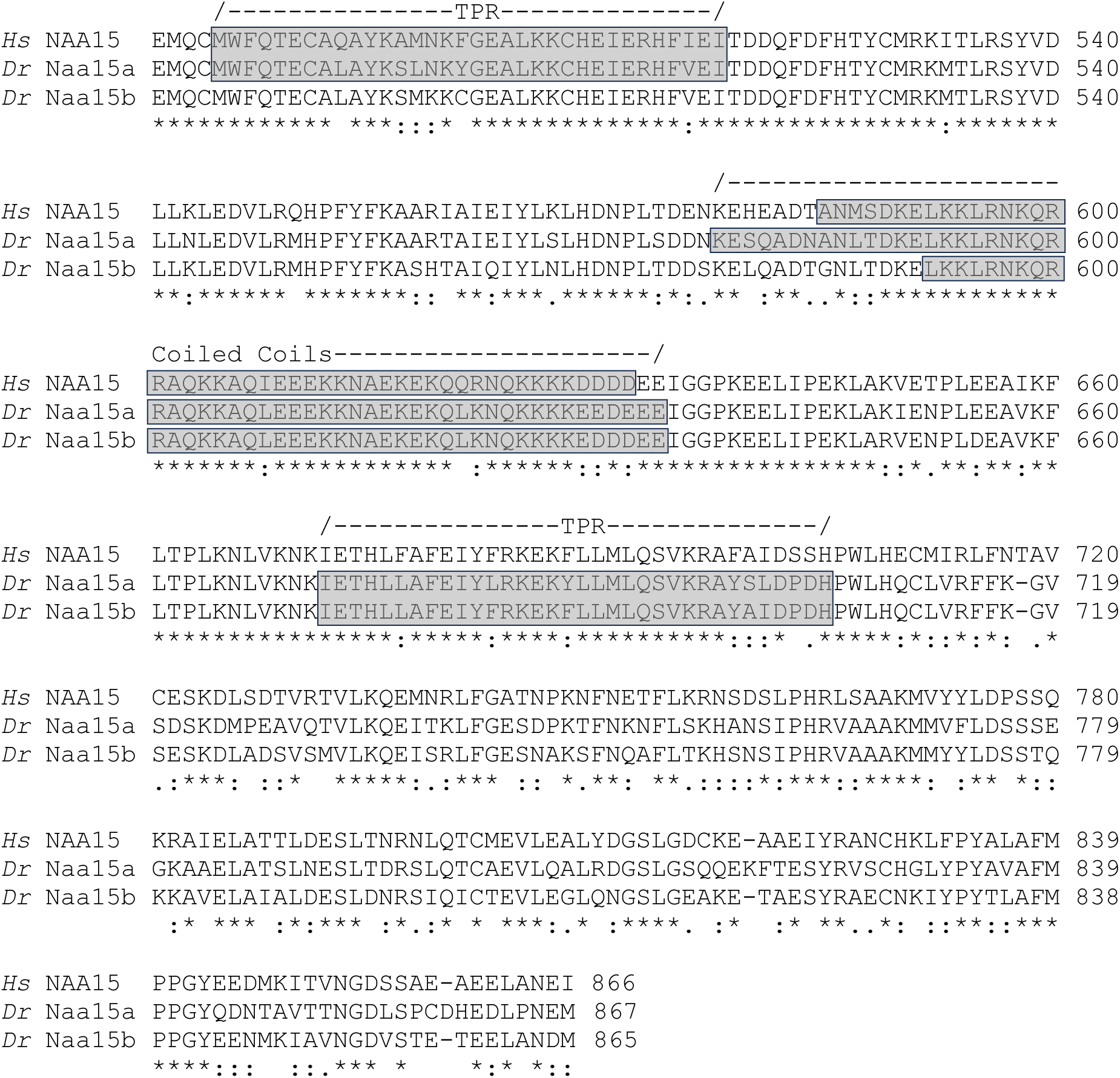
Sequence conservation between human and zebrafish NAA15 proteins. (A) Percent identity matrix for *Homo sapiens* (*Hs*) NAA15, *Danio rerio* (*Dr*) Naa15a, and *Dr* Naa15b. (B) Clustal Omega alignment of *Hs* Naa15, *Dr* Naa15a, and *Dr* Naa15b. Asterisks indicate conserved amino acids, colons highlight partially conserved amino acids or strong similarities in amino acid properties, and periods identify amino acids with weak similarities. Tetratricopeptide repeats (TPR) and coiled-coil domains predicted by InterPro sequence searches are shown.

**Figure S2.**
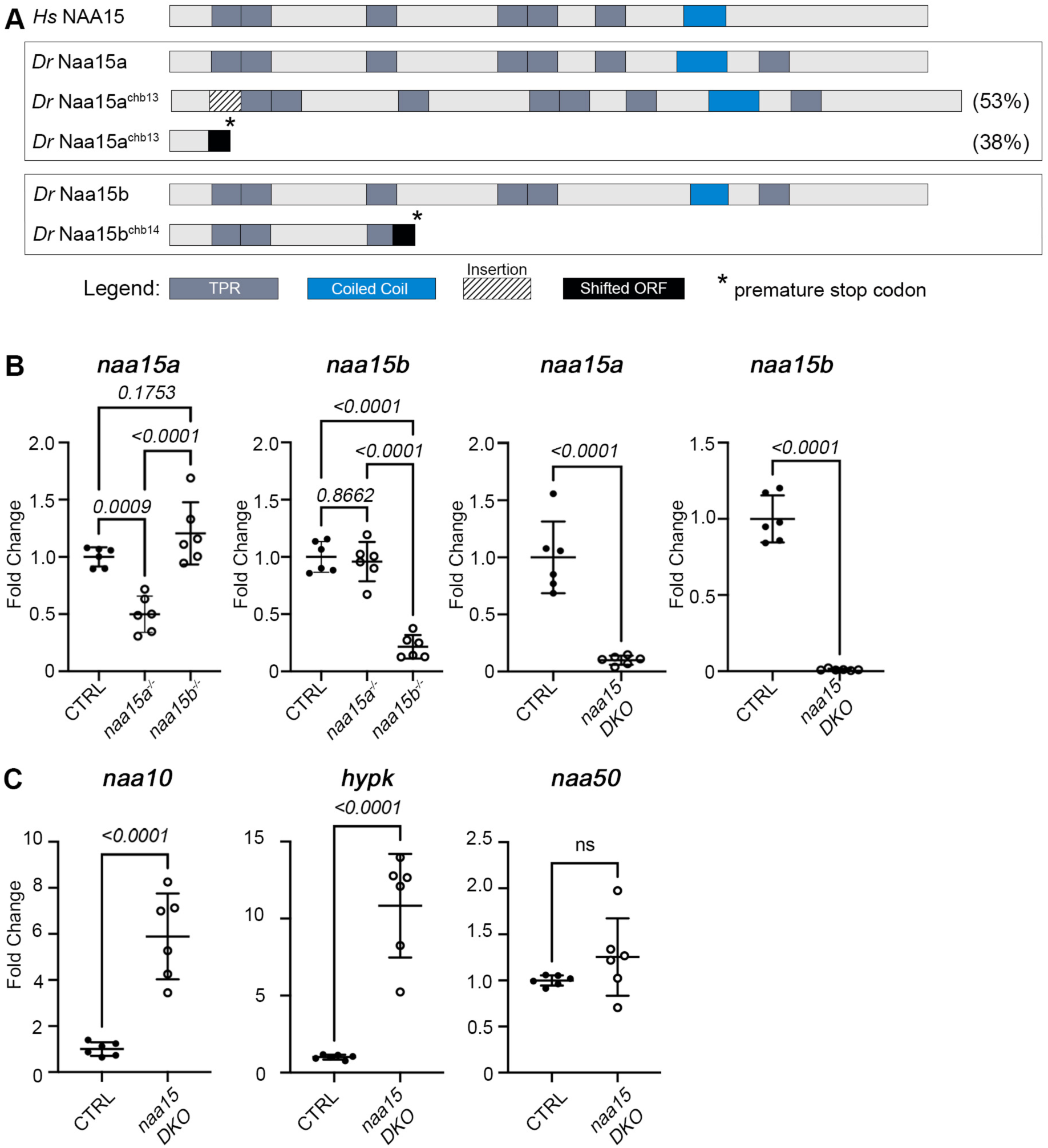
Generation and validation of *naa15*-deficient zebrafish. (A) Schematic diagrams of the domain structures of *Homo sapiens* (*Hs*) NAA15, *Danio rerio* (*Dr*) Naa15a, and *Dr* Naa15b, as well as the predicted protein products of the *naa15a^ch13^* and *naa15b^ch14^* null alleles. (B) Dot plots showing the relative expression levels of *naa15a* and *naa15b* in single-mutant and double-knockout (DKO) animals 4 days post-fertilization (dpf) compared to wild-type control (CTRL) animals. (C) Dot plots showing the relative expression levels of *naa10, naa50,* and *hypk* in DKO animals 4 dpf relative to CTRL animals. n=6 biological replicates per group, each containing 2 or more technical replicates. Statistical significance was determined by ordinary one-way ANOVAs, followed by a Tukey’s multiple comparisons test, for the single-mutant analysis (adjusted p-values are shown) and Student’s t-tests (p-values are shown) for the DKO analysis. Means ± 1SD are shown. Abbr: ns, not significant.

**Figure S3.**
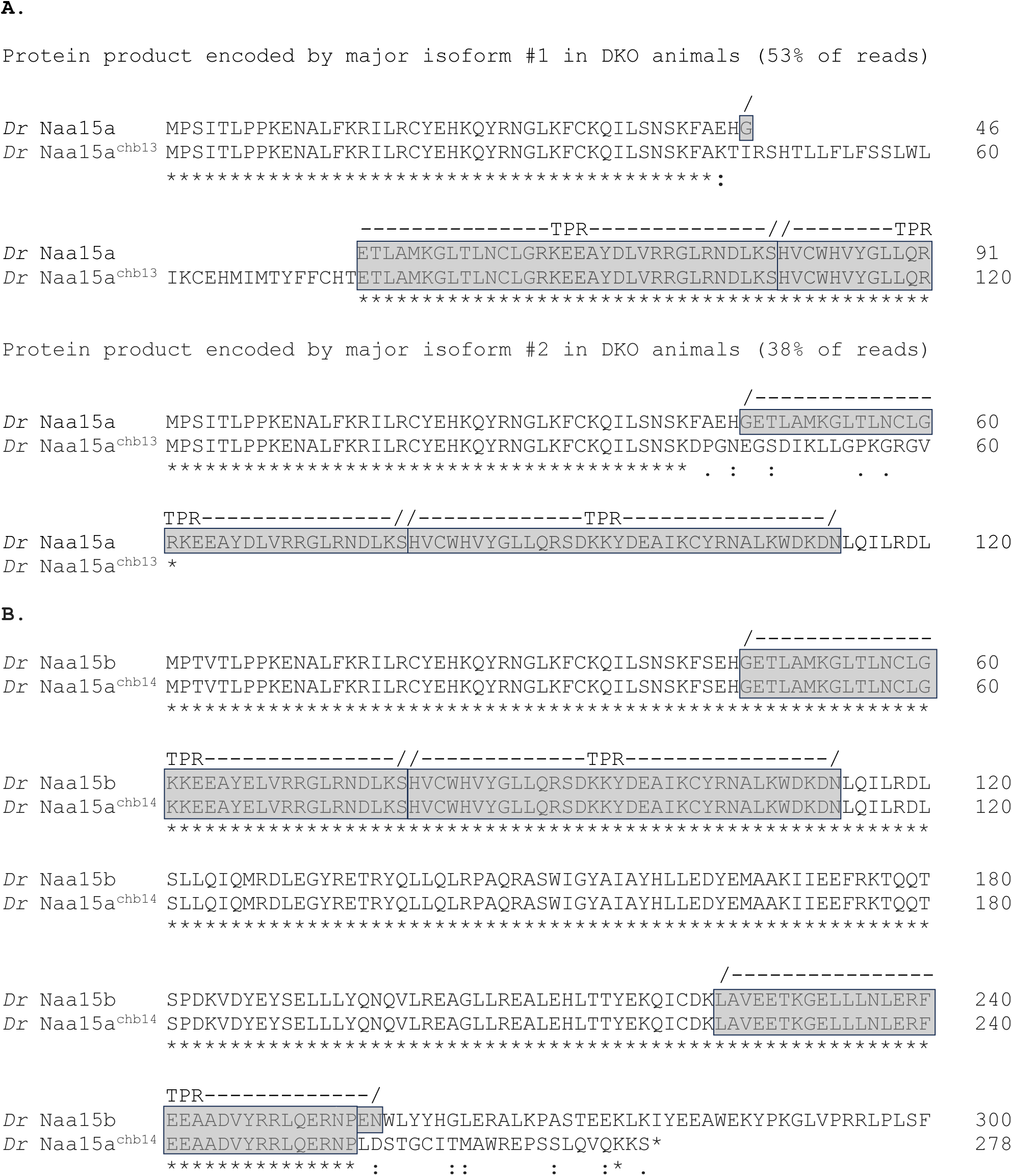
Predicted protein products of *naa15a^chb13^* and *naa15^chb14^* alleles. (A,B) Clustal Omega alignments of wild-type Naa15a (A) and Naa15b (B) *Danio rerio* (*Dr*) proteins and the predicted protein products of the *naa15a^chb13^*(two isoforms; A) and *naa15^chb14^* (B) alleles. The relevant partial WT sequences are shown. Asterisks indicate conserved amino acids, colons highlight partially conserved amino acids or strong similarities in amino acid properties, and periods identify amino acids with weak similarities.

**Figure S4.**
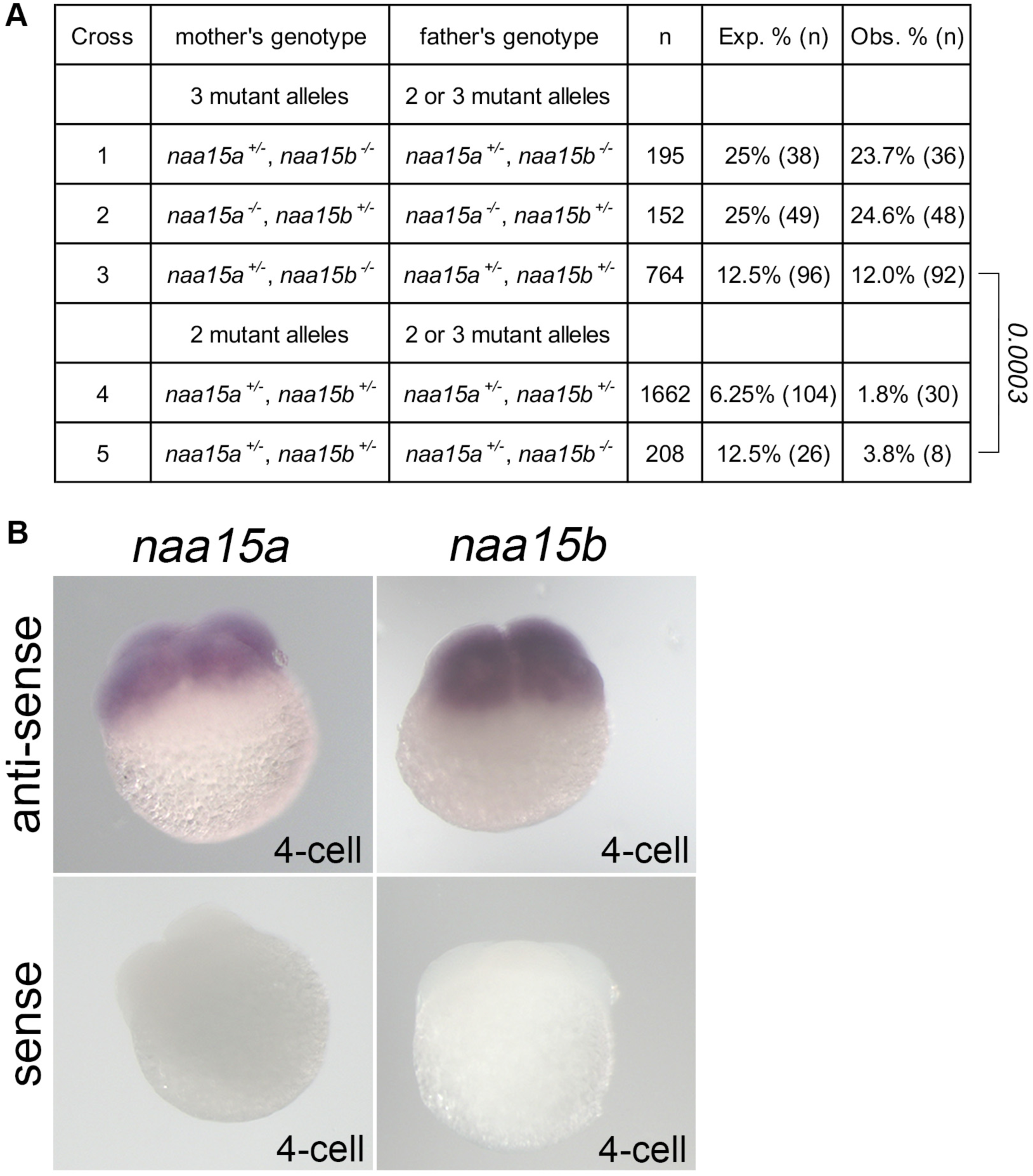
*naa15a* and *naa15b* are maternal-effect genes. (A) Table showing the expected and observed percentages of animals showing cardiac edema on 4 days post-fertilization in clutches from parents with the indicated genotypes. The observed number of phenotypic animals followed a Mendelian inheritance pattern when the mother carried three mutant alleles in either combination (crosses 1-3). However, the observed number was lower than expected, ostensibly due to increased maternal load when the mother was double heterozygous (crosses 4,5). The statistical significance between the outcomes of crosses 3 and 5 was determined by a Fisher’s Exact Test. (B) Representative brightfield images of 4-cell stage wild-type embryos processed for in situ hybridization to detect *naa15a* and *naa15b* messages with anti-sense riboprobes. Also shown are control embryos exposed to sense probes. N=25/25 embryos per group showed similar staining patterns.

**Figure S5.**
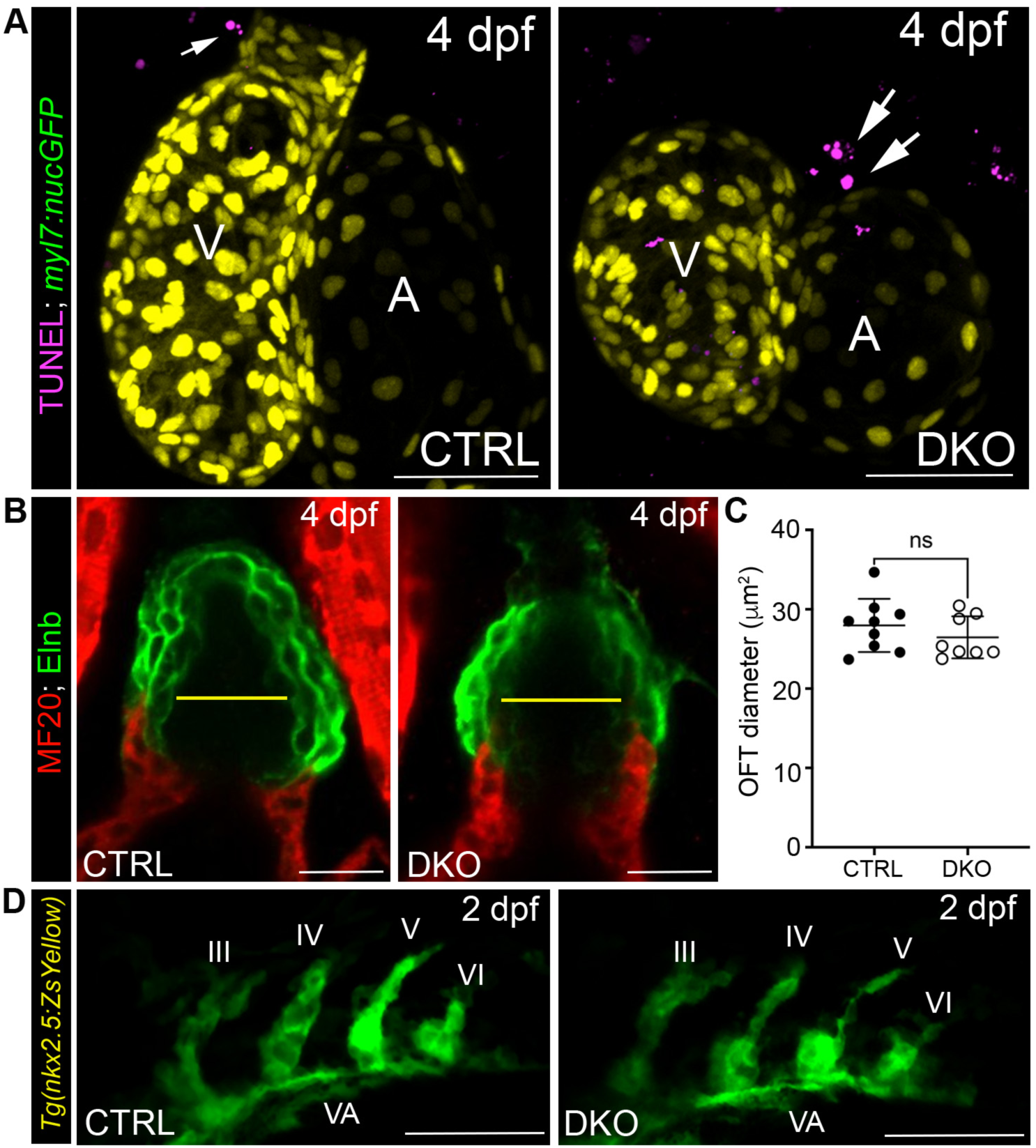
*naa15* deficiency does not induce CM apoptosis or affect morphogenesis of the outflow tract or pharyngeal arch arteries. (A) Representative confocal projections of hearts in 4 days post-fertilization (dpf) control-sibling (CTRL) and double knockout (DKO) animals carrying the *myl7:nucGFP* transgene processed for TUNEL staining. No TUNEL+ CM nuclei were observed in CTRL. (n=7) or DKO hearts (N=13). The arrows highlight TUNEL signals in non-CMs. (B) Single optical sections through the outflow tracts (OFTs) of 4 dpf CTRL and DKO animals double immunostained to detect myocardium (MF20 antibody) and Elastin b (Elnb). (C) Dot plot showing OFT diameters in 4 dpf CTRL (n=9) and DKO (n=8) animals. Statistical significance was determined by a Student’s t-test. Means ± 1SD are shown. ns, not significant. (D) Representative confocal projections of pharyngeal arch (PA) arteries III-VI in live 2 dpf CTRL and DKO animals carrying the *nkx2.5:ZsYellow* transgene. Similar vessel morphologies were observed in CTRL (n=5) and DKO (n=4) animals. Abbr: V, ventricle; A, atrium; VA, ventral aorta. Scale bars=25μM.

**Figure S6.**
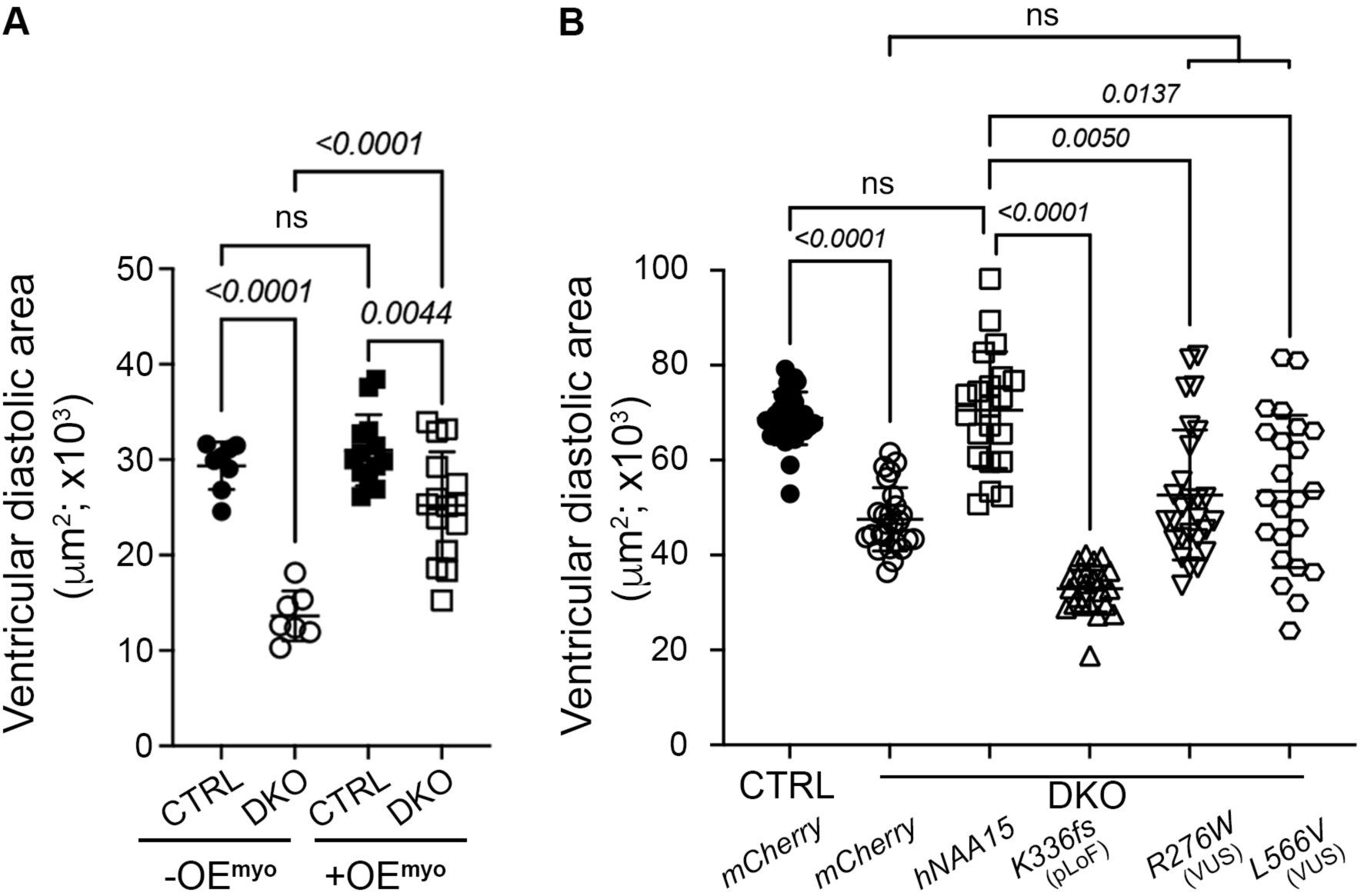
Ventricular size analysis of OE^myo^ and human *NAA15*-expressing larval hearts. (A) Dot plot showing the ventricular end-diastolic areas in 4 dpf control-sibling (CTRL) and double-knockout (DKO) animals without (CTRL, n=8; DKO, n=7) and with (CTRL, n=13; DKO, n=16) overexpression of *naa15a* in the myocardium. (B) Dot plot showing the ventricular end-diastolic areas in 4 dpf CTRL and DKO animals injected at the one-cell stage with control mCherry (CTRL, n=30; DKO, n=27), WT human (h) *NAA15* (n=21), and variant-containing *hNAA15* (k336fs, n=24; R276W, n=27; L566V, n=23) mRNAs. Statistical significance was determined by an ordinary one-way ANOVA (A) or a Kruskal-Wallis Test followed by a Dunn’s multiple comparisons test (B). Means ± 1SD are shown, as are adjusted p-value upper limits. Abbr: pLoF, predicted loss of function; VUS, variant of uncertain significance.

**Figure S7.**
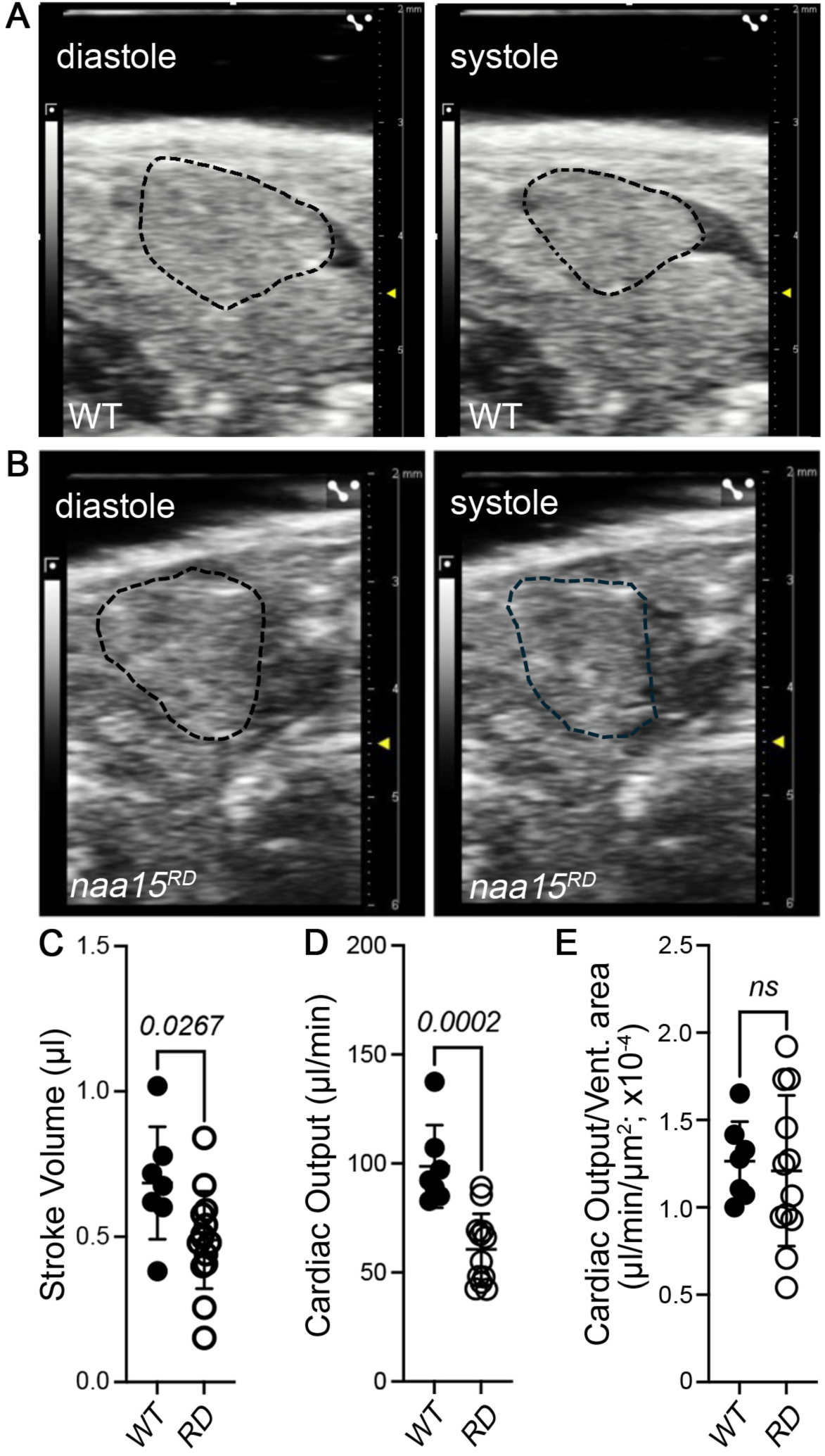
B-mode ultrasound analysis of adult *naa15^RD^* hearts. (A,B) B-mode ultrasound images of 9 months post-fertilization (mpf) WT and *naa15^RD^* ventricles during systole and diastole. The ventricles are outlined by dotted lines. (C,D) Dot plots showing the stroke volumes (C), cardiac outputs (D), and cardiac outputs normalized to ventricular areas of 9 mpf WT (n=7) and *naa15^RD^*(n=12) hearts. Statistical significance was determined with Student’s t-tests. Means ± 1SD are shown.

**Figure S8.**
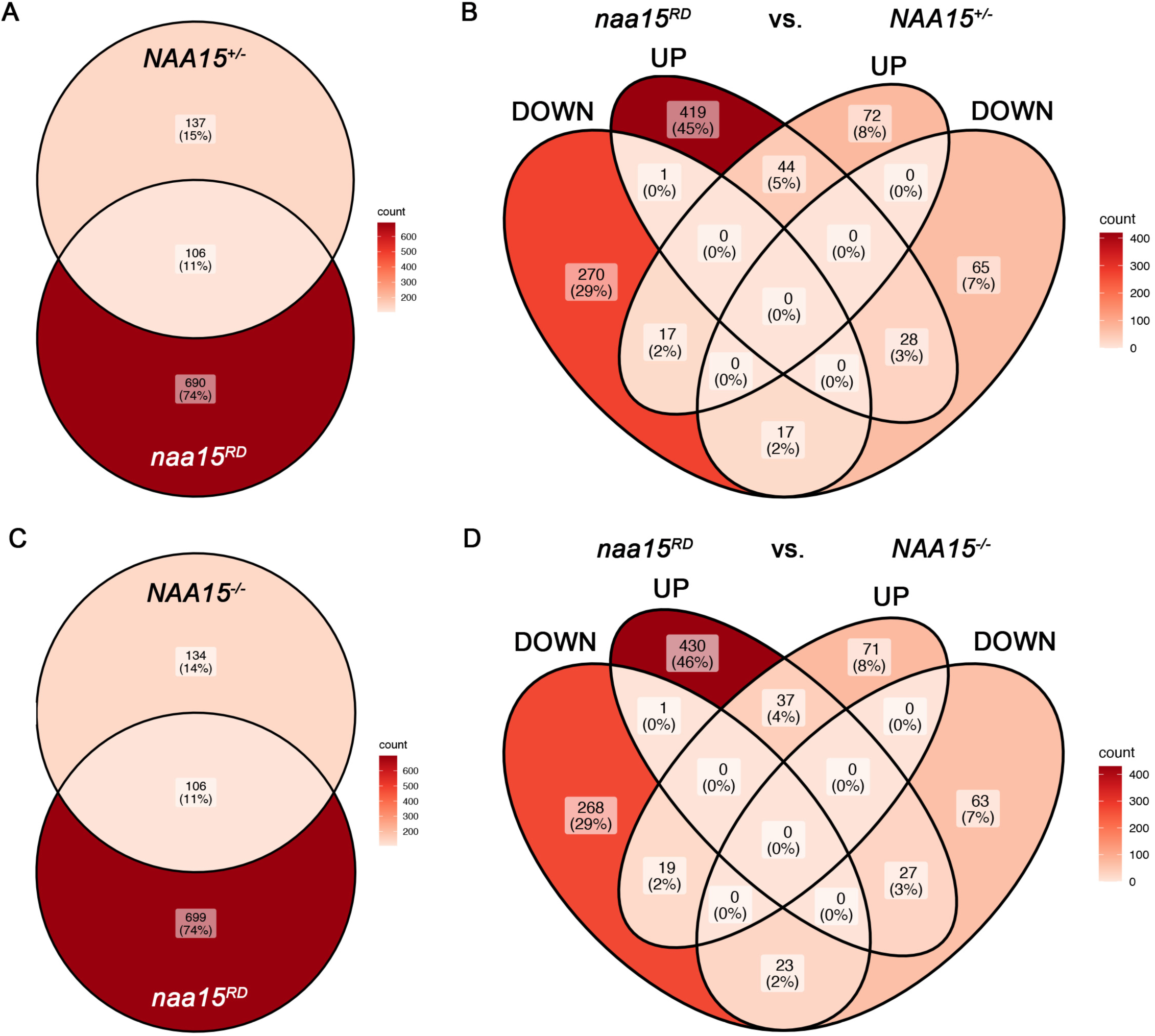
Cross-species comparison of differentially expressed proteins resulting from reduced or absent *NAA15* activity. (A-D) Venn diagrams showing the degrees of overlap between zebrafish proteins differentially expressed in *naa15^RD^* hearts (A-D) and human proteins differentially expressed in *NAA15* heterozygous (*NAA15^+/-^*; A,B) or null (*NAA15^-/-^*; C,D) induced pluripotent stem cells (iPSCs). The numbers and percentages were derived from differential expression data for zebrafish-human protein orthology pairs (Table 5) identified as described (Materials and Methods). The degrees of overlap between datasets with (B,D) or without (A,C) consideration of whether proteins were UP- or DOWN-regulated are presented. Similarities between the dysregulated proteins in zebrafish and human were not statistically significant by Fisher’s Exact tests (Table 5). Zebrafish proteins with two measurements were counted as single proteins when both were significant and moving in the same direction.

## Notes

### Competing Interest Statement

The authors have declared no competing interest.

### Summary of Updates

Updated to include new data and new and revised text.

